# Poly (ADP-ribose) polymerase-1 regulates HIV-1 replication in human CD4+ T cells

**DOI:** 10.1101/2024.06.11.598467

**Authors:** Zachary S. Martinez, Denisse A. Gutierrez, Carlos Valenzuela, Chang-Soo Seong, Manuel Llano

**Author notes:** Corresponding author. Address correspondence to: Manuel Llano. Department of Biological Sciences. University of Texas at El Paso. 500 West University Ave. El Paso, TX 79968. USA.

## Abstract

The cellular enzyme poly (ADP-ribose) polymerase-1 (PARP-1) regulates multiple processes that are potentially implicated in HIV-1 infection. However, the role of PARP-1 in HIV-1 infection remains controversial, with reports indicating or excluding that PARP-1 influence early steps of the HIV-1 life cycle. Most of these studies have been conducted with Vesicular Stomatitis virus Glycoprotein G (VSV-G)-pseudotyped, single-round infection HIV-1; limiting our understanding of the role of PARP-1 in HIV-1 replication. Therefore, we evaluated the effect of PARP-1 deficiency or inhibition in HIV-1 replication in human CD4+ T cells. Our data showed that PARP-1 knockout increased viral replication in SUP-T1 cells. Similarly, a PARP-1 inhibitor that targets PARP-1 DNA-binding activity enhanced HIV-1 replication. In contrast, inhibitors affecting the catalytic activity of the enzyme were inactive. In correspondence with the pharmacological studies, mutagenesis analysis indicated that the DNA-binding domain was required for the PARP-1 anti-HIV-1 activity, but the poly-ADP-ribosylation activity was dispensable. Our results also demonstrated that PARP-1 acts at the production phase of the viral life cycle since HIV-1 produced in cells lacking PARP-1 was more infectious than control viruses. The effect of PARP-1 on HIV-1 infectivity required Env, as PARP-1 deficiency or inhibition did not modify the infectivity of Env-deleted, VSV-G-pseudotyped HIV-1. Furthermore, virion-associated Env was more abundant in sucrose cushion-purified virions produced in cells lacking the enzyme. However, PARP-1 did not affect Env expression or processing in the producer cells. In summary, our data indicate that PARP-1 antagonism enhances HIV-1 infectivity and increases levels of virion-associated Env.

**Importance:** Different cellular processes counteract viral replication. A better understanding of these interfering mechanisms will enhance our ability to control viral infections. We have discovered a novel, antagonist effect of the cellular enzyme poly (ADP-ribose) polymerase-1 (PARP-1) in HIV-1 replication. Our data indicate that PARP-1 deficiency or inhibition augment HIV-1 infectivity in human CD4+ T cells, the main HIV-1 target cell *in vivo*. Analysis of the mechanism of action suggested that PARP-1 antagonism increases in the virus the amounts of the viral protein mediating viral entry to the target cells. These findings identify for the first time PARP-1 as a host factor that regulates HIV-1 infectivity, and could be relevant to better understand HIV-1 transmission and to facilitate vaccine development.

## INTRODUCTION

Poly (ADP-ribose) polymerase-1 (PARP-1) is a key cellular enzyme implicated in multiple cellular processes, including DNA repair and regulation of transcription (1, 2). PARP-1 is the most enzymatically active member of the PARP family and promotes the transfer of ADP ribose molecules from NAD+ to acceptor proteins or to an existing poly (ADP-ribose) chain (3). This post-translational modification modifies the function of target proteins by altering their subcellular localization, molecular interactions, and enzymatic activities. In addition, catalytic-independent functions have been demonstrated to mediate PARP-1 biological activities (2–6).

PARP-1 is abundant in cells that *in vivo* support HIV-1 replication and is implicated in different cellular processes that could be important for HIV-1 infection. However, the role of this enzyme in HIV-1 infection is controversial. Several reports have indicated that PARP-1 affects early steps in the viral life cycle in human cells, including reverse transcription (7), HIV-1 DNA integration (8, 9), and gene expression (7, 10, 11) (12, 13); while others exclude a fundamental role of PARP-1 in HIV-1 infection (14–16). However, most of these studies have been conducted with Vesicular Stomatitis Virus Glycoprotein G (VSV-G)-pseudotyped, single-round infection HIV-1, hindering our understanding of the role of PARP-1 in HIV-1 replication. Therefore, we evaluated the effect of PARP-1 deficiency or inhibition in HIV-1 replication in human CD4+ T cells. We have found that PARP-1 deficiency or inhibition enhances HIV-1 infectivity in an Env-dependent manner. Our data also exclude a fundamental role of PARP-1 in the early steps for the HIV-1 life cycle in human CD4 T cells. The anti-HIV-1 function of PARP-1 requires the DNA-binding but not the poly-ADP-ribosylation activity of the enzyme. Suggesting a potential mechanism, we found that Env was more abundant in virions produced in cells lacking PARP-1 than in control viruses. However, PARP-1 deficiency did not alter the production or processing of Env in the producer cells. Therefore, we propose that PARP-1 could contribute to the notorious low abundance of Env in HIV-1 virions.

## RESULTS

### PARP-1 deficiency enhances HIV-1 replication in human CD4+ T cells

HIV-1 replication was evaluated in PARP-1-knockout clones (KO) derived from the human CD4 T cell line SUP-T1 and their wild-type PARP-1-backcompleted (BC) counterparts (**Fig. 1a**). KO and BC cells were infected with HIV-1_NL4-3_ and viral replication was followed for 10 - 12 days by HIV-1 p24 ELISA. In these experiments, we observed that HIV-1 replicated more robustly in cells lacking PARP-1 than in the backcomplemented cells (**Fig. 1b**). As expected, all the cells analyzed in **Fig. 1b** expressed almost identical CD4 and CXCR4 levels by FACS analysis (**Fig. 1c**).

**Figure 1.**
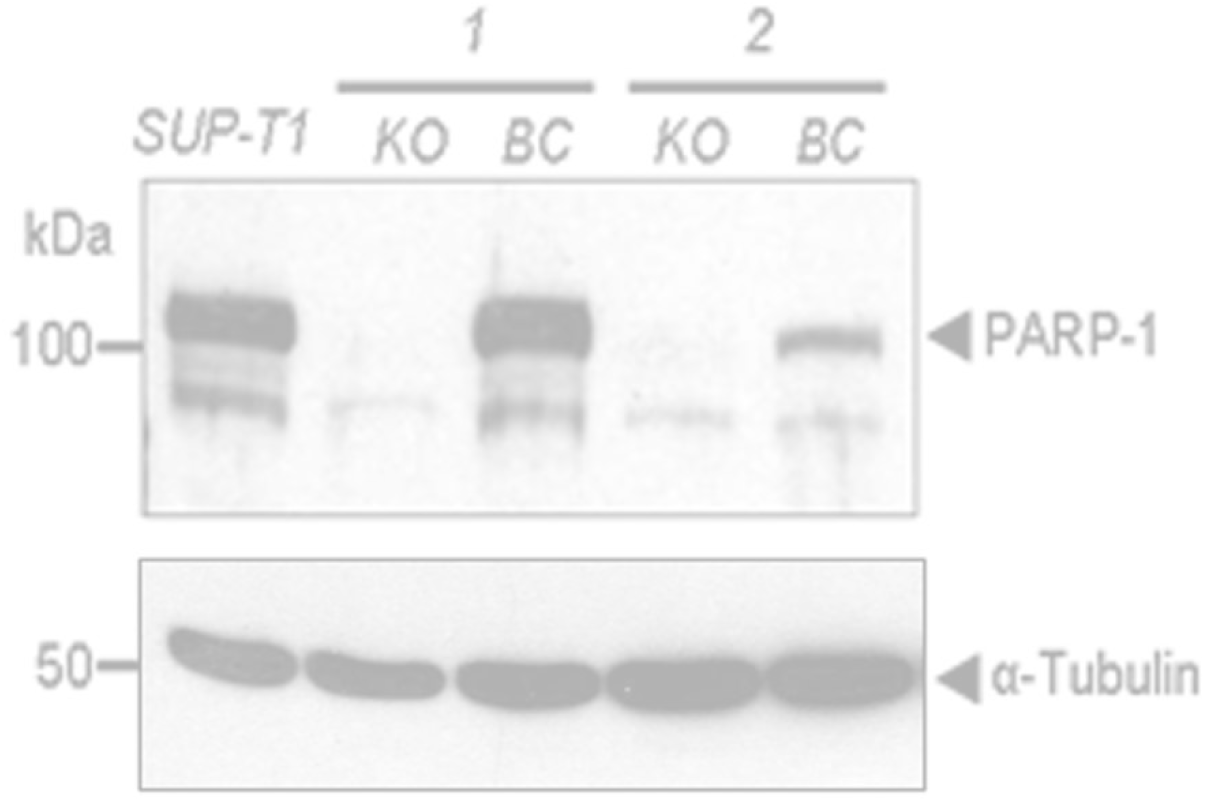
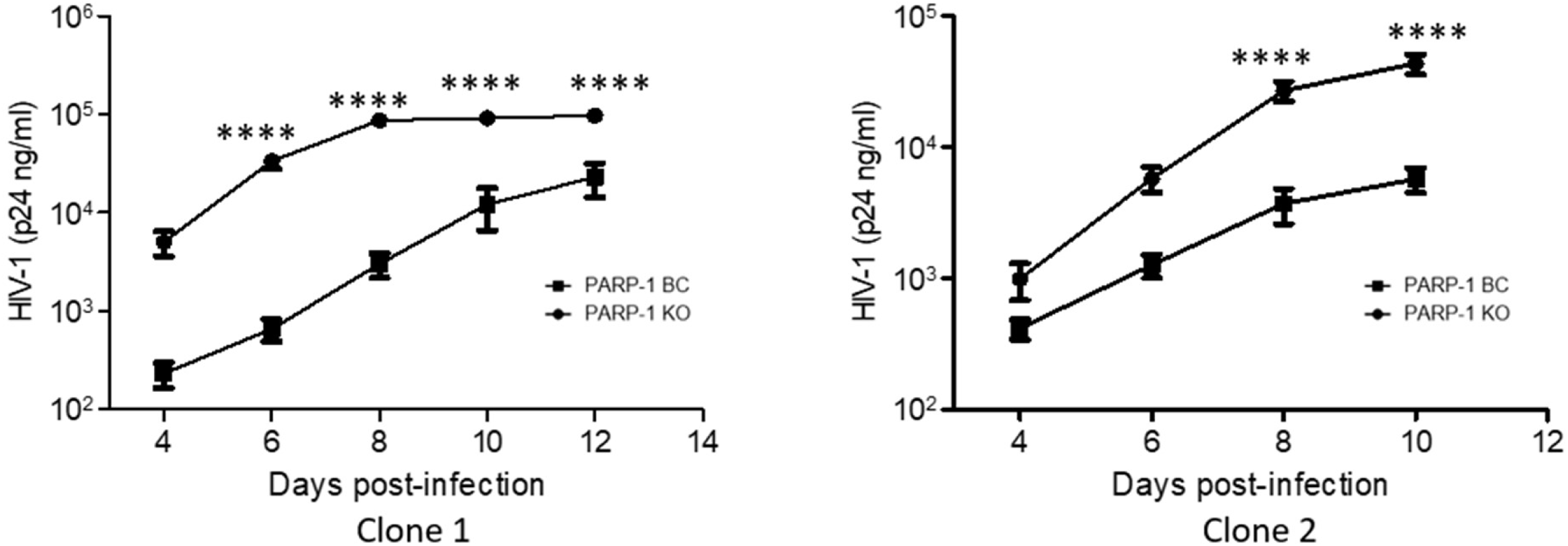
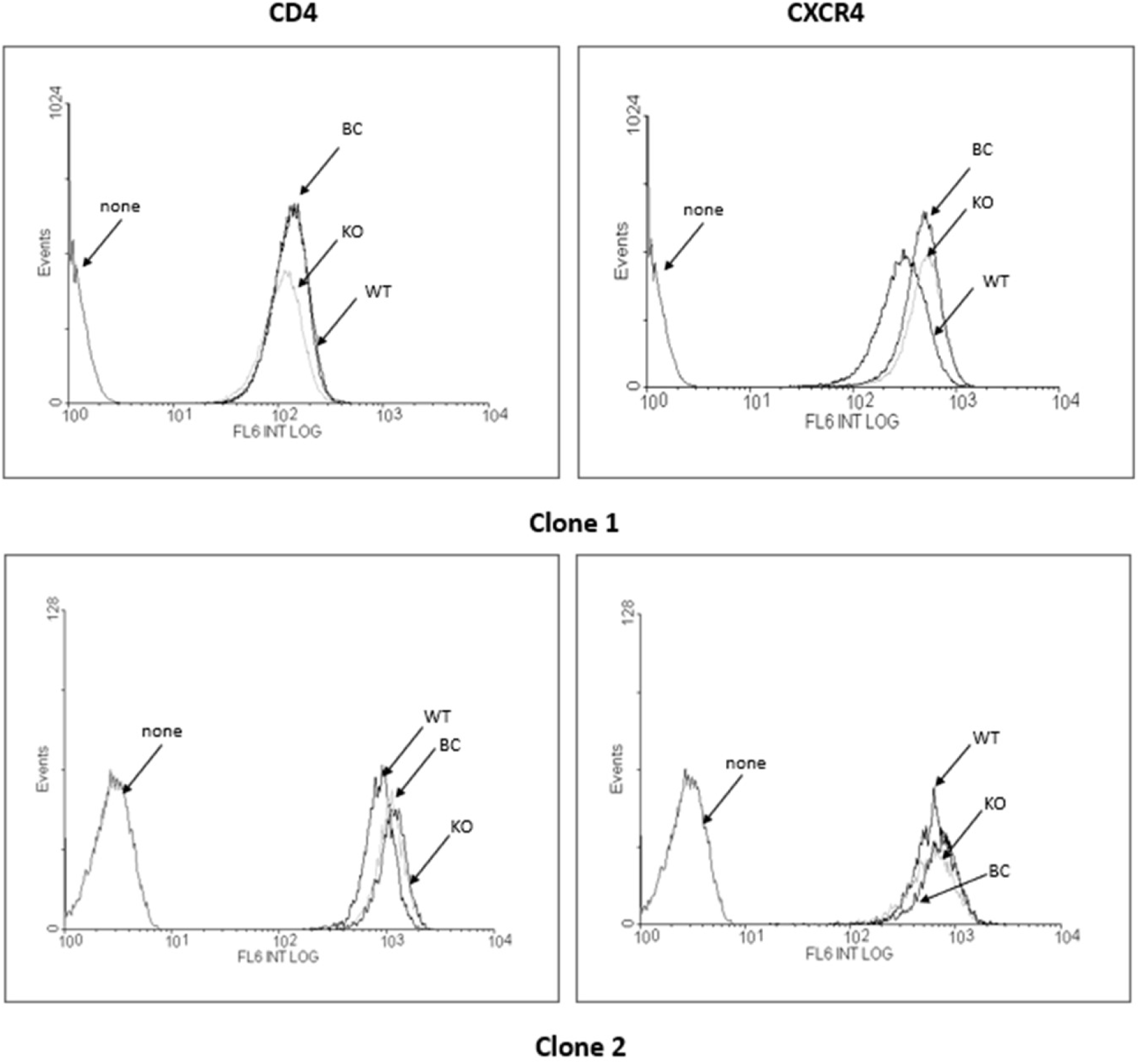
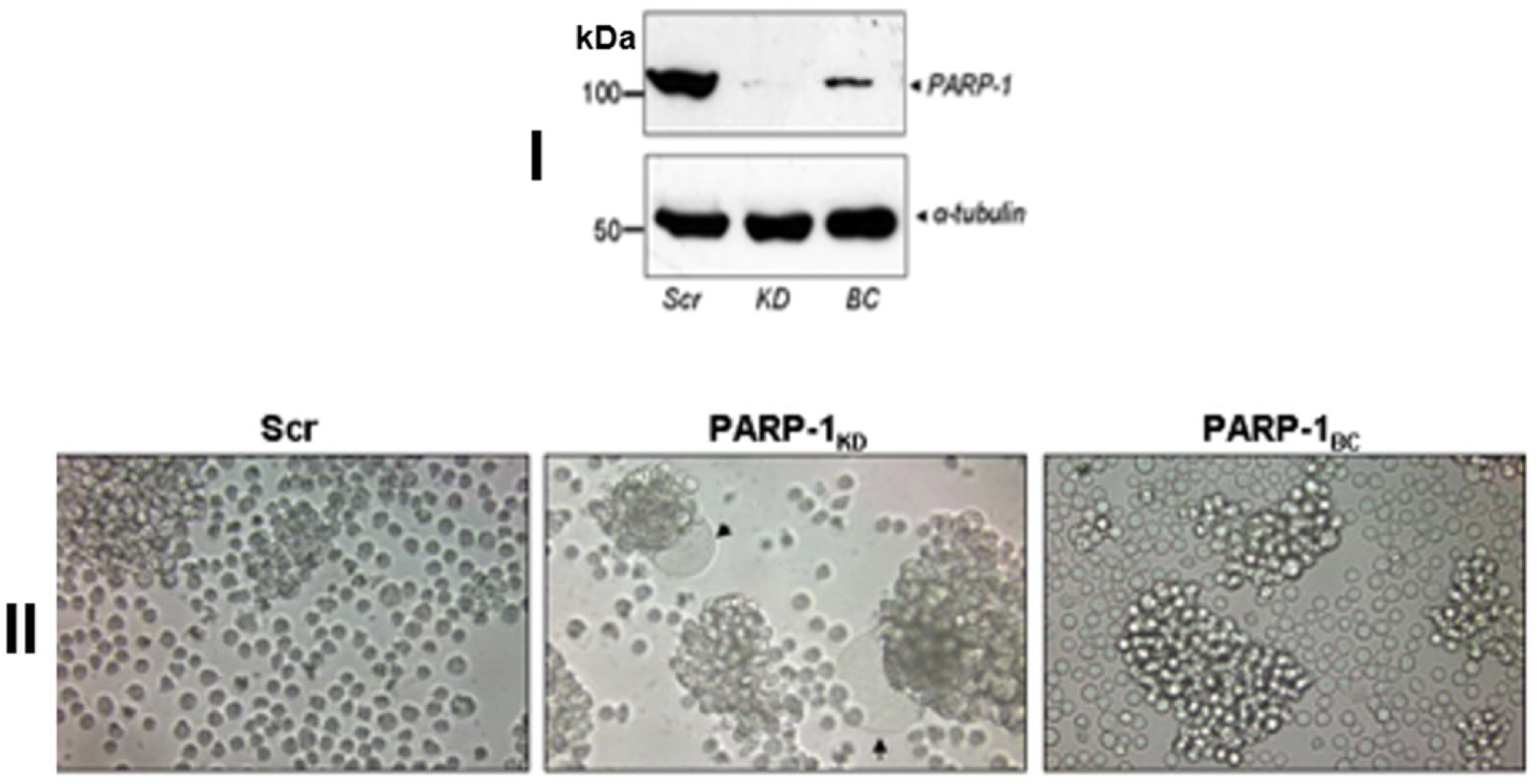
PARP-1 deficiency in human CD4 T cells enhances HIV-1 replication. (**a**) PARP-1 expression in two PARP-1 knockout (KO) SUP-T1-derived clones and their corresponding backcomplemented (BC) lines. (**b**) HIV-1_NL4-3_ replication in PARP-1 KO and BC clones studied in panel **a**. HIV-1 p24 was determined by ELISA. Results correspond to one experiment done in triplicate and are representative of more than five independent experiments. Statistical analysis was performed by a multiple paired t-test with the Bonferroni-Dunn method, p >0.0001 (****). Non-significant differences (p </= 0.05) are not indicated. (**c**) Cell surface expression of CD4 and CXCR4 in the clones represented in (**a**) and (**b**). WT refers to parental SUP-T1 cells. (**d-I**) PARP-1 expression in shRNA-PARP-1 knockdown (KD) SUP-T1-derived cells, the corresponding backcomplemented (BC) cell line, and a SUP-T1-derived cell line expressing a scrambled shRNA sequence (Scr). (**d-II**) HIV-1_NL4-3_-induced syncytia in cells in panel **d-I** at eight days post-infection. Results are representative of more than three independent experiments.

An important consideration in these experiments was the amount of virus used. We noticed that the phenotype reported in figure **1b** was routinely obtained across several viral preps with low doses of HIV-1 p24 (between 2.4 and 15 ng / 0.25 x 10^6^ cells). Control SUP-T1 cells infected with these low viral doses, showed syncytia and produced between 0.1-1ug/ml of p24 by day 7-9 post-infection. Higher HIV-1 doses produced a more robust viral replication in control SUP-T1 cells, but the enhancing effect of PARP-1 on HIV-1 replication was less noticeable. Since a similar dose-dependency was reported for the enhancing effect of Env-induced CD4/CXCR4 signaling on HIV-1 replication (17), we evaluated the effect of PARP-1 deficiency in virus-induced syncytia, an Env-dependent event. SUP-T1 cells rendered PARP-1 knockdown (KD) by stable expression of an shRNA (18), and their wild-type PARP-1-BC counterpart (**Fig. 1d-I**) were infected with HIV-1_NL4-3_ and syncytia formation evaluated by microscopy analysis. Larger and more abundant virus-induced syncytia were observed in PARP-1-KD than in the -BC cells (**Fig. 1d-II**), suggesting an involvement of Env in the PARP-1 phenotype.

### PARP-1 deficiency enhances viral infectivity in an Env-dependent manner at a late step of the viral life cycle

To further verify the Env-dependency on the PARP-1 effect on HIV-1 infection, we evaluated the susceptibility of PARP-1-KO and -BC cells to the infection with a VSV-G-pseudotyped, *env*-deleted single-round infection HIV-1 (Hluc). This reporter virus derives from HIV-1_NL4-3_ and expresses LTR-driven luciferase from the *nef* slot (19, 20). Luciferase levels measured in PARP-1-KO and -BC cells 4 days after Hluc infection indicated that PARP-1-KO cells were three-fold less susceptible than -BC cells (**Fig. 2a**), in contrast to the observed enhancing effect of PARP-1 deficiency on wild-type HIV-1 replication.

**Figure 2.**
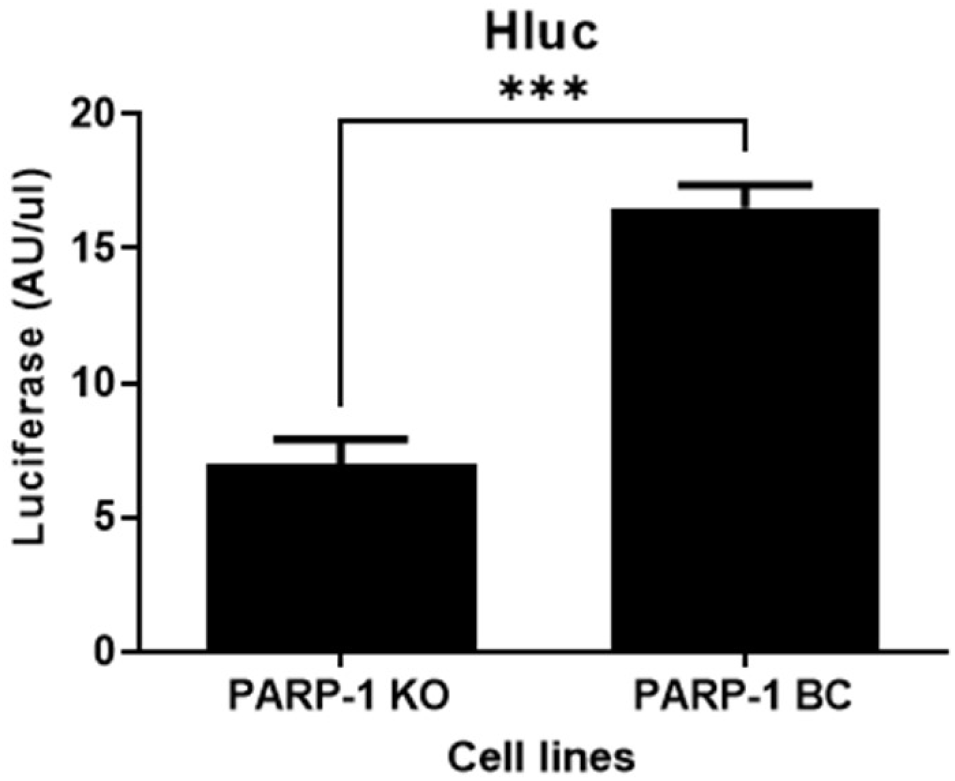
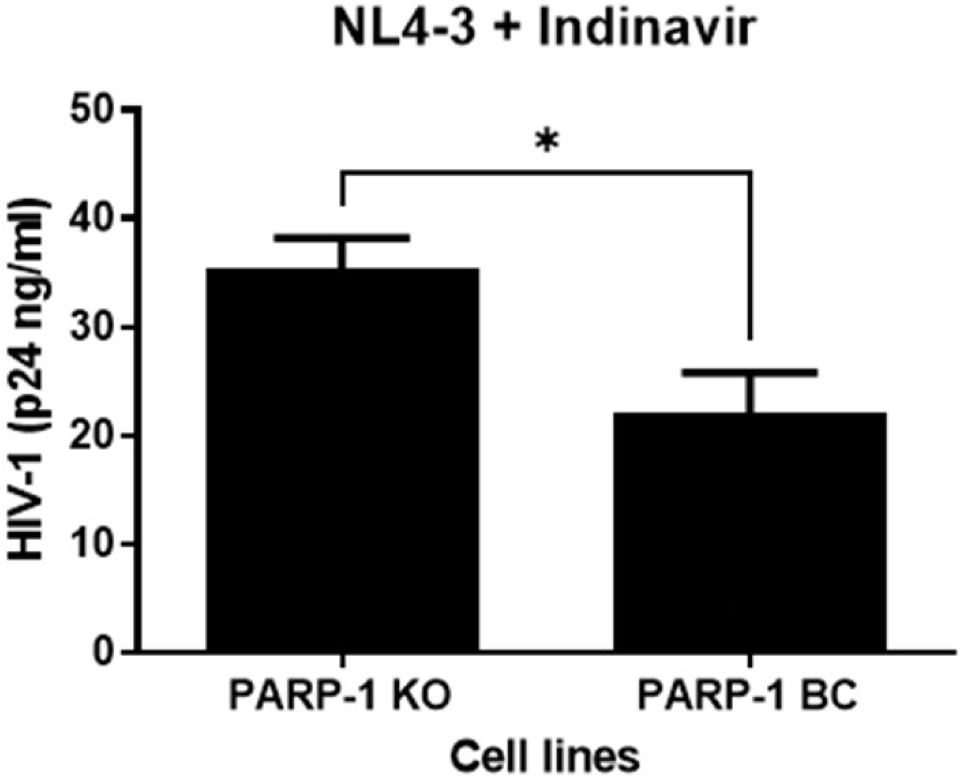
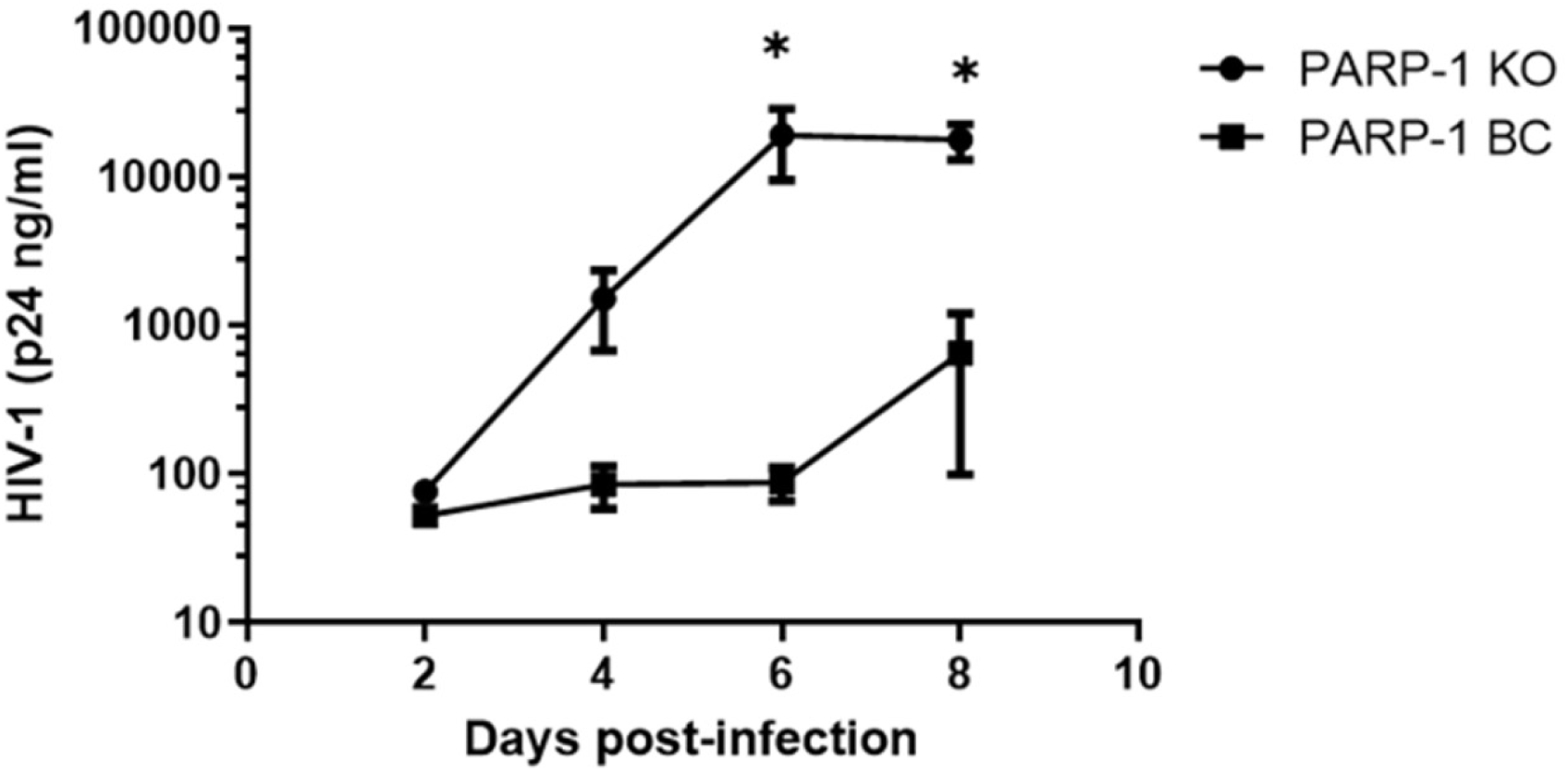
PARP-1 deficiency enhances HIV-1 replication in an Env-dependent manner. (**a**) Infection of PARP-1 KO and BC cells with a VSV-G-pseudotyped HIV-1 lacking Env, Vpr, and Nef that expresses LTR-driven luciferase (Hluc). Results correspond to one experiment conducted in triplicate and are representative of more than three independent experiments. (**b**) Infection of PARP-1 KO and BC cells with HIV-1_NL4-3_ in the presence of Indinavir (100 nM). (**c**) Indinavir was removed in cells evaluated in panel **b**, and viral replication was followed by HIV-1 p24 ELISA. Results in panels **b** and **c** correspond to the same triplicate experiment and are representative of two other independent experiments. In (**a**) and (**b**) a two-tailed t-test statistical analysis was performed. In (**c**), a 2-way ANOVA with a Sidak post-hoc test was used. p > 0.05 (*) and p >0.001 (***). Non-significant differences (p </= 0.05) are not indicated.

To determine whether the enhancing effect of PARP-1 deficiency on viral replication was at an early or late step of the viral life cycle, we infected PARP-1-KO and -BC cells with HIV-1_NL4-3_ in the presence of Indinavir (100 nM) to restrict infection to a single-round. HIV-1 p24 levels measured two days later indicated minor differences (∼ 1.6-fold) in HIV-1_NL4-3_ infectivity in PARP-1 KO and BC cells (**Fig. 2b**). However, after removing the indinavir blockage, HIV-1 replicated significantly more vigorously in PARP-1-KO than -BC cells (**Fig. 2c**), as previously observed. These results suggested that PARP-1 affects late rather than early steps of the HIV-1 life cycle.

To evaluate the effect of PARP-1 on the late steps of the HIV-1 life cycle, we produced Hluc and HIV-1_NL4-3_ in HEK293T PARP-1-KO and -BC cells (**Fig. 3a**) by transfecting the corresponding expression plasmids, and HIV-1 p24-normalized viruses were used to infect the reporter cell line TZM-bl. Reporter cells infected with HIV-1_NL4-3_ were treated with indinavir (100 nM) or raltegravir (60 nM) at the time of viral challenge to limit the infection to a single round. Data in **Fig. 3b** indicated that HIV-1_NL4-3_ (**b-I** and **b-II**), but no Hluc (**b-III**), produced in the absence of PARP-1 were more infectious, suggesting that PARP-1 enhances viral infectivity during the production phase in an Env-dependent manner. Moreover, p24-normalized HIV-1_NL4-3_ produced in HEK293T PARP-1-KO was more infectious in SUP-T1 cells than viruses produced in HEK293T PARP-1-BC cells (**Fig. 3c-I**). This effect that was not observed with Hluc produced in HEK293T PARP-1-KO and -BC cells (**Fig. 3c-II**).

**Figure 3.**
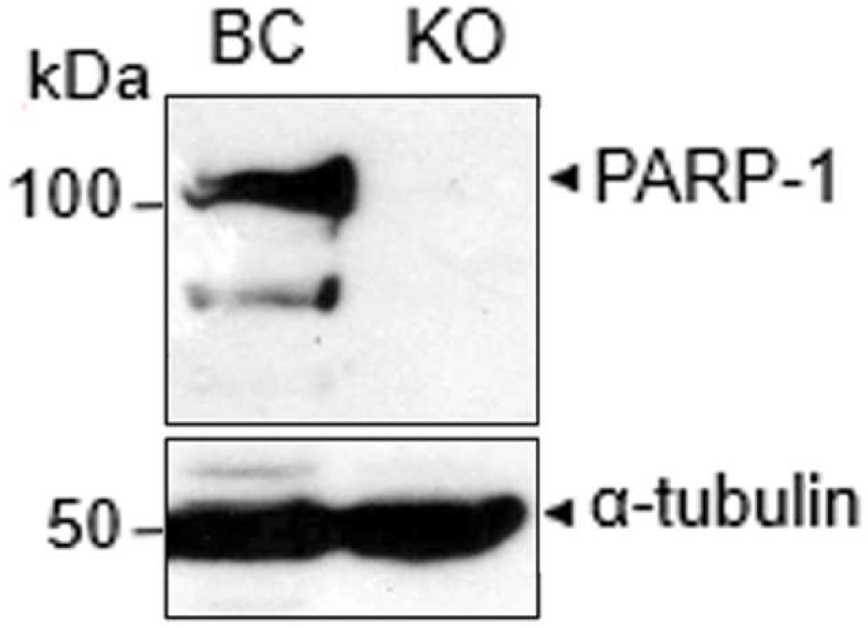
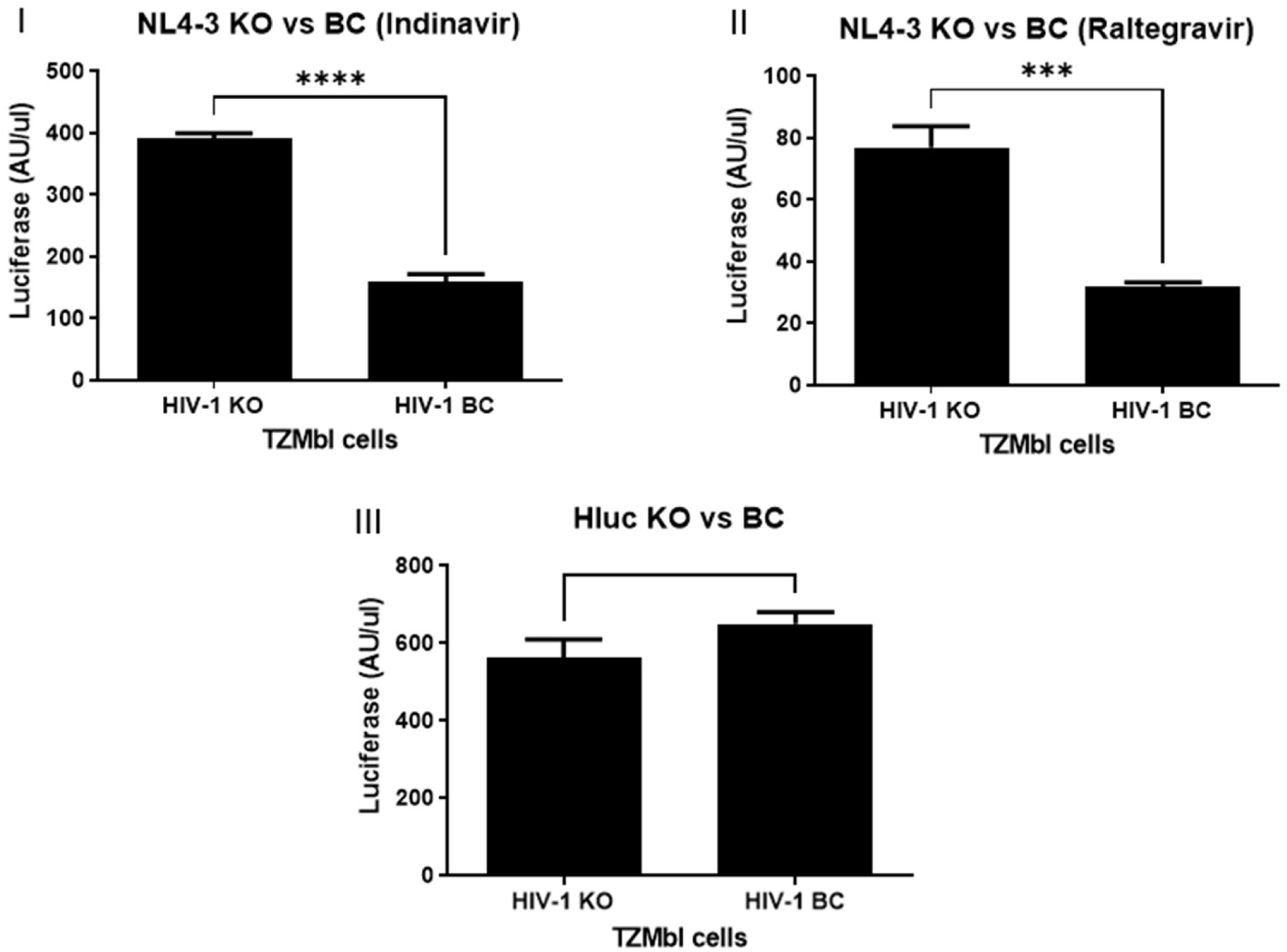
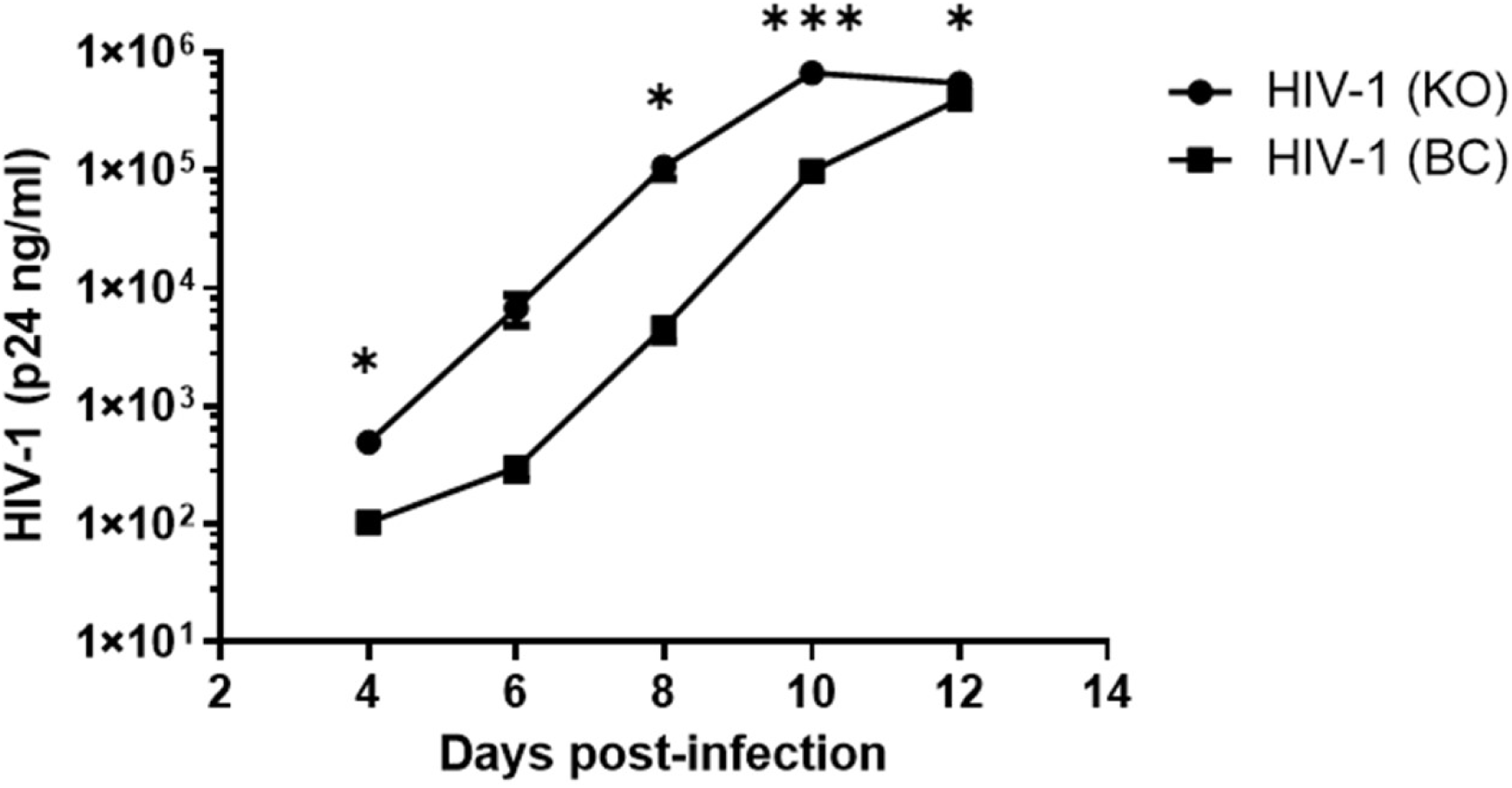
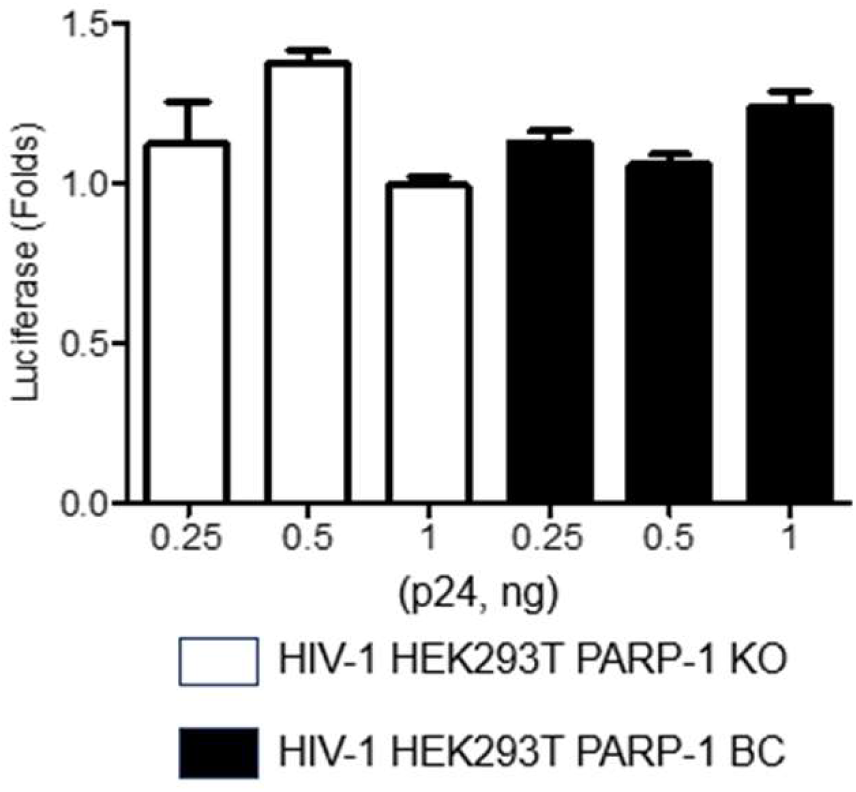
PARP-1 deficiency enhances HIV-1 infectivity at a late step of the viral life cycle. (**a**) PARP-1 expression in PARP-1 knockout (KO) HEK293T-derived cells and their corresponding backcomplemented (BC) lines.(**b**). TZM-bl cells were infected with HIV-1_NL4-3_ (**b-I** and **b-II**) or Hluc (**b-III**) produced in KO and BC HEK293T-derived cells. HIV-1_NL4-3_ infections were carried out in the presence of indinavir (100 nM) or raltegravir (60 nM). (**c**) SUP-T1 cells were infected with HIV-1_NL4-3_ (**I**) or Hluc (**II**) produced in KO and BC HEK293T-derived cells. Several doses of Hluc were used and luciferase values were normalized to those in cells infected with 1 ng of HIV-1 HEK293T PARP-1 KO. Results in **b** and **c** are from one triplicate experiment and are representative of three independent experiments. In (**b**), statistical analysis was performed by a two-tailed t-test, and in (**c-I**) by a multiple paired t-test with the Bonferroni-Dunn method. p > 0.05 (*), >0.001 (***), and >0.0001 (****). Non-significant differences (p </= 0.05) are not indicated.

Furthermore, we evaluated the effect of PARP-1 deficiency on the infectivity of other single-round infection HIV-1 viruses. VSV-G-pseudotyped *env* and *env*/*vpr*/*nef* deleted versions of HIV-1_NL4-3_ were produced in HEK293T PARP-1-KO and -BC cells, p24-normalized, and used to infected SUP-T1 cells. Infection was determined four days later, by measuring the levels of HIV-1 p24 in the supernatant of the infected cells. SUP-T1 cells infected with VSV-G pseudotyped HIV-1_NL4-3ΔEnv_ or HIV-1_NL4-3ΔEnv/Vpr/Nef_ viruses produced in HEK293T PARP-1-KO produced 83 +/- 6.5 % and 71.8 +/- 5.9 %, respectively, of the p24 levels obtained with these viruses produced in HEK293T PARP-1-BC cells. These results further demonstrated that PARP-1 deficiency did not enhance the infectivity of VSV-G pseudotyped HIV-1.

### PARP-1 pharmacological antagonism enhances HIV-1 replication

To better characterize the mechanism implicated, we determined the effect of PARP-1 pharmacological interference on HIV-1 replication. Most of the PARP inhibitors are nicotinamide mimetics that inhibit the enzymatic activity of PARP-1, and several other members of the PARP family by competing with NAD+ for binding to a pocket in the catalytic domain (21). In contrast, only one PARP-1 inhibitor, 5-iodo-6-amino-1,2-benzopyrone (INH2BP), that ejects zinc from the zinc-finger domains is commercially available (22, 23). INH2BP affects only PARP-1 within the PARP family and primarily impairs the DNA binding ability of PARP-1 that is required for activation of the enzymatic activity (21, 24, 25). Therefore, we determined the effect of the nicotinamide mimetic inhibitor NU1025 and the zinc ejector INH2BP on HIV-1 replication.

SUP-T1 cells were infected with HIV-1_NL4-3_ in the presence of DMSO, NU1025 (0.5 mM) or INH2BP (50 uM), and 18 hrs later the cells were washed and cultured for 2 weeks in the absence of these compounds. HIV-1 replication was followed by microscopy evaluation of syncytia formation. As early as day 8 post-infection, syncytia was very evident in cells treated with the zinc-ejector INH2BP, but not with DMSO or the nicotinamide mimetic inhibitor NU1025 (**Fig. 4a-I**), and these differences in cytopathic effect correlated with higher levels of HIV-1 p24 in their cell culture supernatant (**Fig. 4a-II**). Similarly, another nicotinamide mimetic inhibitor, *3*-Aminobenzamide (3-ABA, 5 mM), also failed to enhance HIV-1 replication (Data not shown).

**Figure 4.**
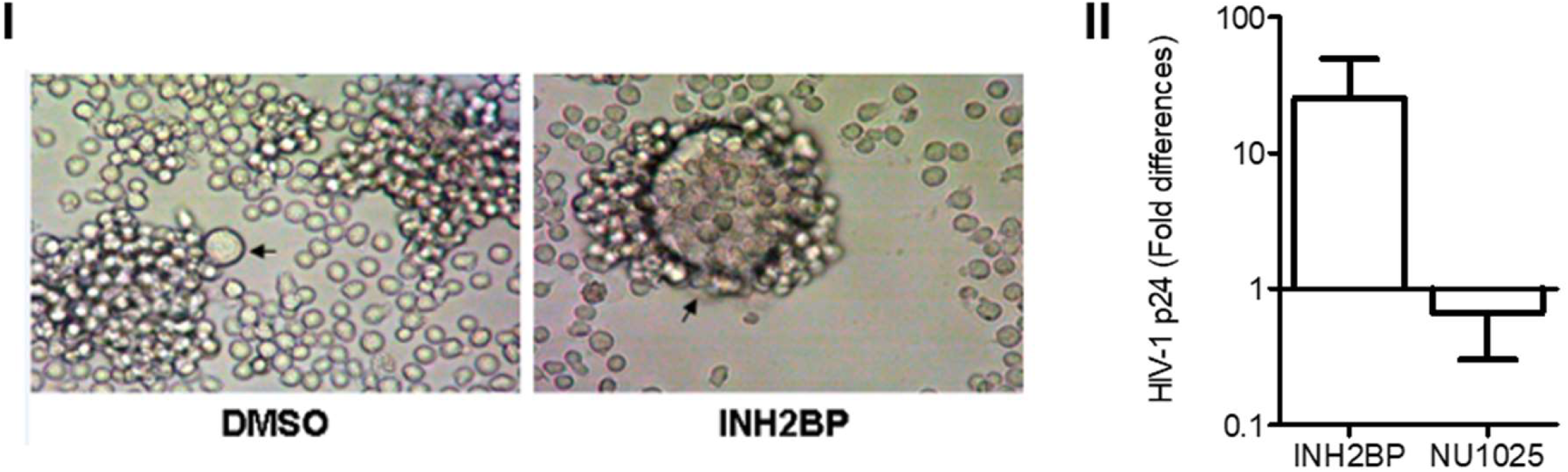
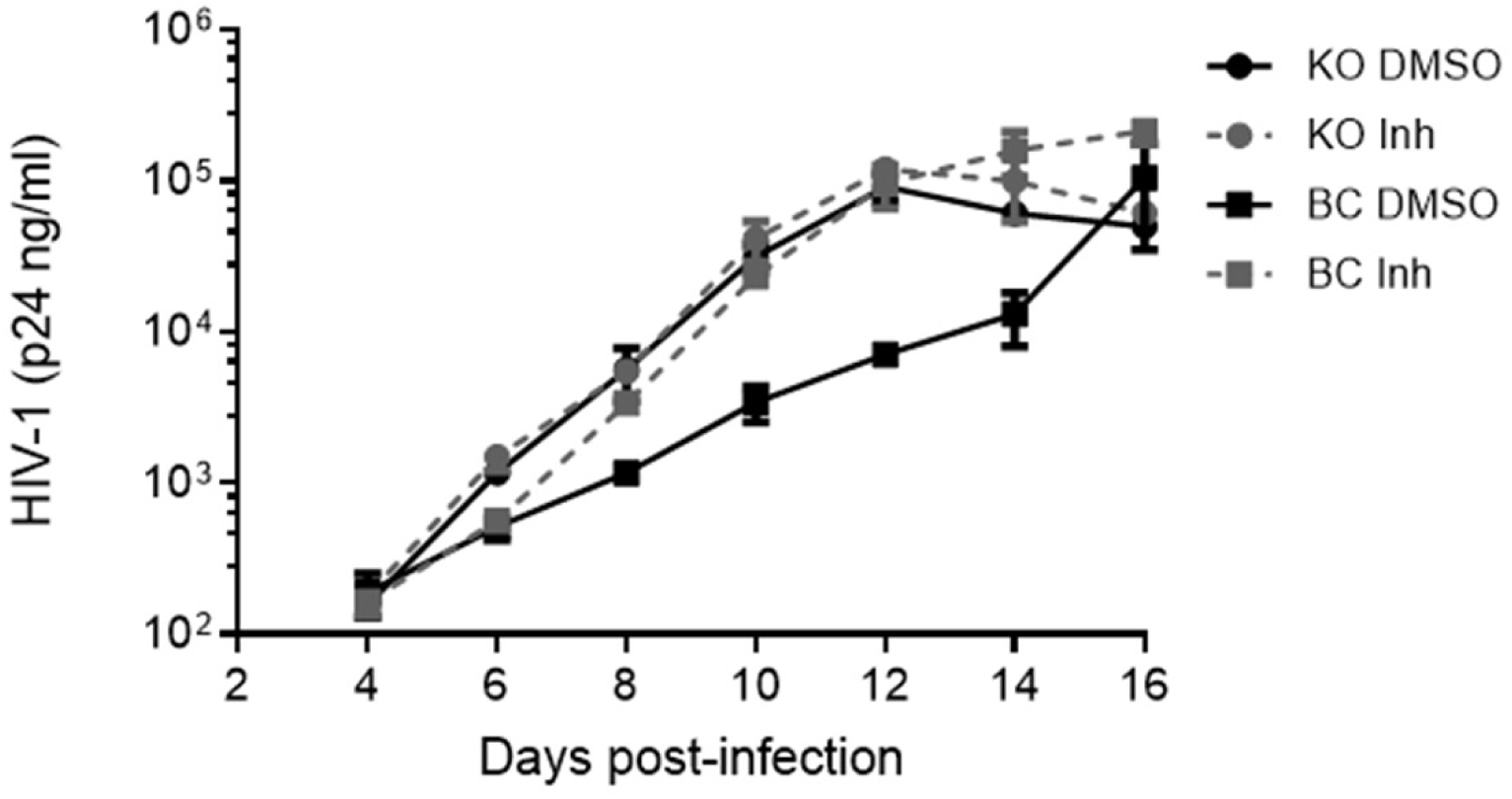
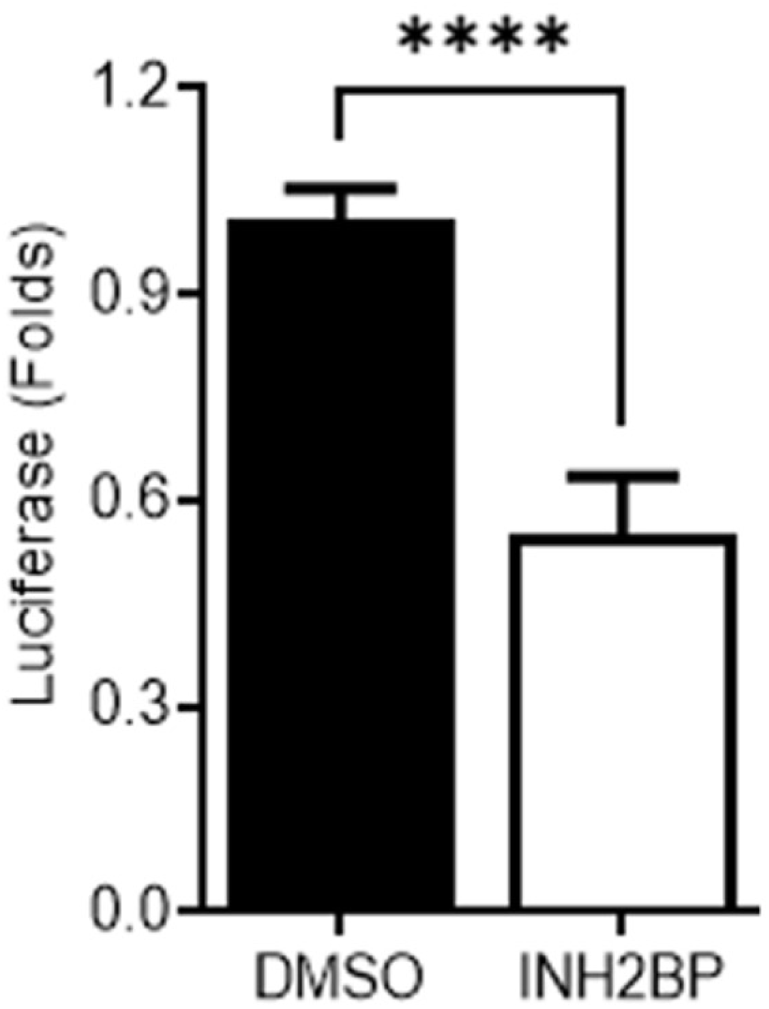
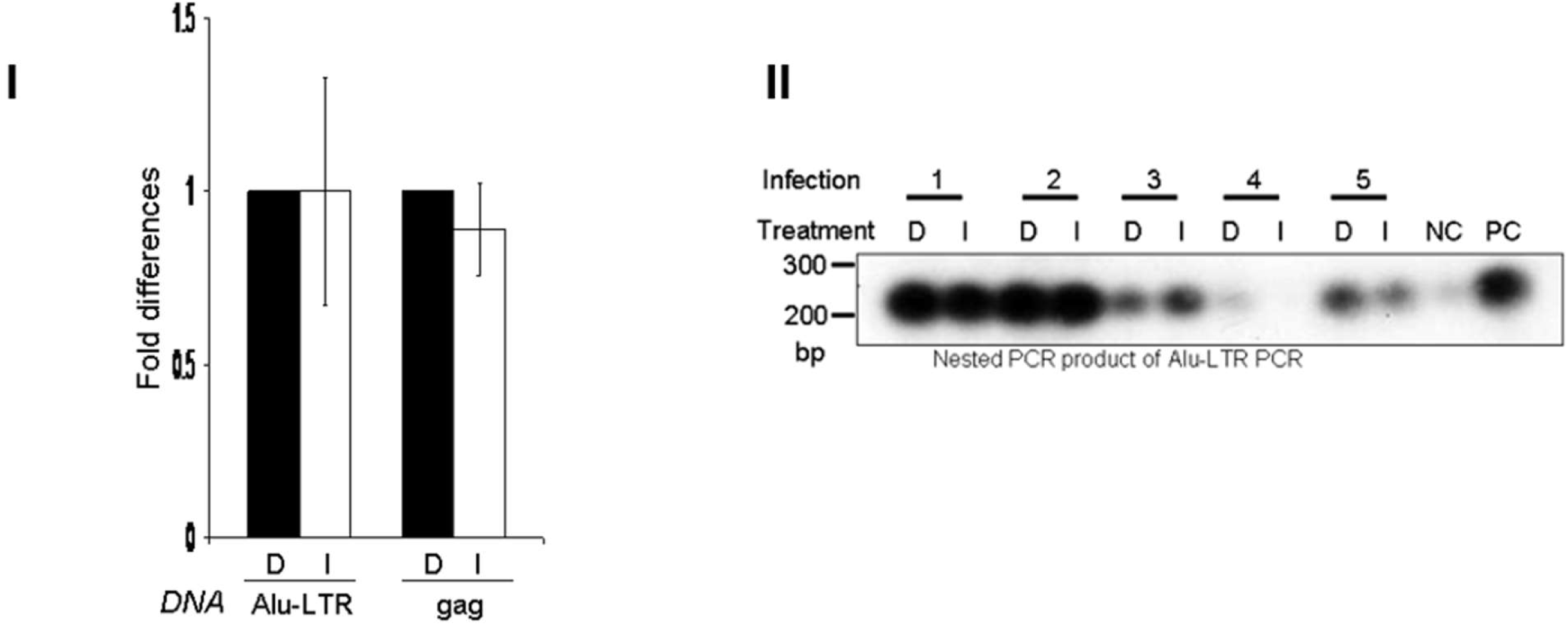
PARP-1 inhibition enhances HIV-1 infectivity in an Env-dependent manner at a late step of the viral life cycle. SUP-T1 cells were infected with HIV-1_NL4-3_ in the presence of NU1025 (0.5 mM), INH2BP (50uM), or DMSO, and viral replication was followed by microscopy evaluation of syncytia formation. (**a-I**) HIV-1-induced syncytia as observed at day 8 post-infection. (**a-II**) HIV-1 p24 production detected by ELISA at day 8 post-infection in cells that are represented in panel (**a-I**). Results correspond to one triplicate experiment and represent more than ten independent experiments. (**b**) SUP-T1-derived PARP-1 KO and BC cells were infected with HIV-1_NL4-3_ in the presence of DMSO or INH2BP (50uM), and viral replication was followed by HIV-1 p24 production by ELISA. Results from one triplicate experiment, representative of two independent experiments, are shown. (**c**) Luciferase levels four days after infection of SUP-T1 cells with Hluc in the presence of DMSO or INH2BP (50uM). Results are from one experiment performed in triplicate and are representative of more than five independent experiments. (**d-I**) Quantification of Alu-LTR junctions and total HIV-1 cDNA (*gag*) in genomic DNA extracted from SUP-T1 cells infected with HIV-1_NL4-3_ in the presence of DMSO or INH2BP (50uM). Results are from an experiment done in triplicate and is representative of more than five independent experiments. (**d-II**) Genomic DNA from Hluc-infected SUP-T1 cells evaluated in panel **d-I** that derived from five independent experiments were analyzed by Southern blot with a radiolabeled probe hybridizing to the HIV-1 LTR. Experimental controls were genomic DNA from uninfected (NC, negative control) and HIV-1 infected (PC positive control) cells. In (**c**) and (**d-I**) statistical analysis was performed by a two-tailed t-test, p >0.0001 (****). Non-significant differences (p </= 0.05) are not indicated.

To correlate the effects of the PARP-1 inhibitors on HIV-1 replication with the enzymatic activity of PARP-1, we measured the cellular levels of poly-ADP-ribose (PAR) by ELISA in SUP-T1 cells treated with DMSO, NU1025, 3-ABA, and INH2BP. We noticed that 15 min after treatment, these inhibitors reduced PAR levels by 95%, as compared to DMSO-treated cells. This level of inhibition persisted at twenty-four hrs post-treatment in cells exposed to NU1025- and 3-ABA but it decayed to 59.1 +/- 1.2 % in cells treated with INH2BP. These findings suggested that Poly ADP-ribosylation activity does not influence HIV-1 replication.

To exclude cell toxicity as the cause for the inactivity of the nicotinamide mimetics inhibitors, we determined cell viability in the treated cells by measuring cellular ATP. Compared to DMSO-treated cells (considered 100% of cell viability), cellular ATP levels were not affected by 24 hrs of treatment with 3-ABA, (5 mM, 134 +/- 10% viability), NU1025 (0.5 mM, 135% +/- 8% viability), or INH2BP (50 uM, 97% +/- 3% viability), demonstrating that these compounds were non-toxic at the doses used. Therefore, analysis of the effect of PARP-1 inhibitors on HIV-1 replication suggests that the DNA binding activity of PARP-1, rather than the enzymatic activity, is implicated in the anti-HIV-1 activity of this protein.

To evaluate the PARP-1-specificity of the effect of INH2BP on HIV-1 replication, excluding the role of other potential targets of this compound, we infected PARP-1-KO and -BC cells in the presence of INH2BP or DMSO and followed HIV-1 replication by p24 ELISA (**Fig. 4b**). Importantly, INH2BP only enhanced viral replication in BC cells, demonstrating that PARP-1 was required for its effect on HIV-1 replication.

### PARP-1 inhibition enhances HIV-1 infection at a post-integration step

We also interrogated the effect of INH2BP on the infectivity of Hluc. SUP-T1 cells were infected with Hluc in the presence of DMSO or INH2BP (50 uM), and luciferase activity was determined 4 days later. In contrast to the enhancing effect of INH2BP on HIV-1_NL4-3_ replication, this compound decreased Hluc infection by two-folds (**Fig. 4c**), indicating Env-dependency of the promoting effect of INH2BP on viral infection.

The lack of effect of PARP-1 deficiency or pharmacological antagonism on the infection of single-round viruses indicates that PARP-1 does not impact the early steps of the viral life cycle. To verify this mechanism further, we defined the effect of PARP-1 inhibition on proviral formation. SUP-T1 cells were infected with different amounts of HIV-1_NL4-3_ in the presence of INH2BP or DMSO. Indinavir (100 nM) was added to these cultures for the entire duration of the experiment to limit infections to a single-round. DNA was extracted from the infected cells at day 4 post-infection, Alu-LTR junctions and total HIV cDNA (gag DNA) were measured by real-time PCR as described in (19). The values for these HIV-1-derived PCR products were normalized to the levels of mitochondrial DNA detected in these same samples. As expected, INH2BP failed to augment the levels of provirus (**Fig. 4d-I**). Furthermore, levels of integrated HIV-1 genome were determined by Southern blot analysis of genomic DNA extracted from the infected SUP-T1 cells that were characterized in figure **4d-I**. A LTR ^32^P-radiolabeled PCR product was used as a probe. In correspondence with the Alu-LTR data, INH2BP failed to augment the levels of provirus (**Fig. 4d-II**). Similar results were obtained when total HIV cDNA was determined by quantitative PCR using primers binding to *gag* (Data not shown). These findings further support a reported negligible role of PARP-1 in HIV-1 DNA integration (14, 16, 18, 26), and are aligned with the lack of effect of PARP-1 deficiency or pharmacological antagonism on the infectivity of single-round infection, VSV-G pseudotyped HIV-1, showed above.

### The DNA-binding but no the poly ADP-ribosylation activity of PARP-1 mediates the anti-HIV-1 effect

INH2BP ejects zinc from zinc fingers (ZnF) in PARP-1. This enzyme possesses three ZnFs, all located in the N-terminus of the protein, ZnF1 and ZnF2 are structurally similar and form the DNA-binding domain of PARP-1, whereas ZnF3 regulates the enzymatic activity but is not part of the DNA-binding domain (27–31). Therefore, to define which PARP-1 function mediates the anti-HIV-1 activity, we replicated HIV-1_NL4-3_ in PARP-1-KO SUP-T1 cells backcomplemented with PARP-1 mutants lacking each of the three ZnF motifs (ΔZnF1, ΔZnF2, and ΔZnF3) (**Fig. 5a-I**). Importantly, the deletion of ZnF1 or ZnF2, but no of ZnF3, caused a significant decrease in the anti-viral activity of PARP-1, indicating that the DNA-binding activity of PARP-1 was required for the anti-HIV-1 activity (**Fig. 5b-I-III**).

**Figure 5.**
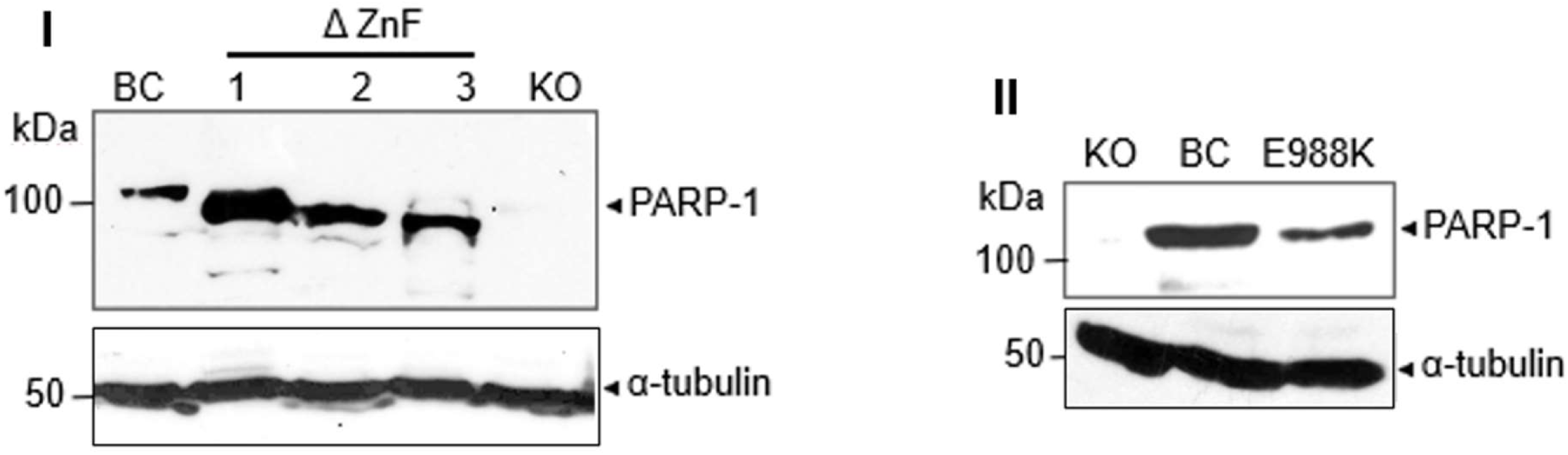
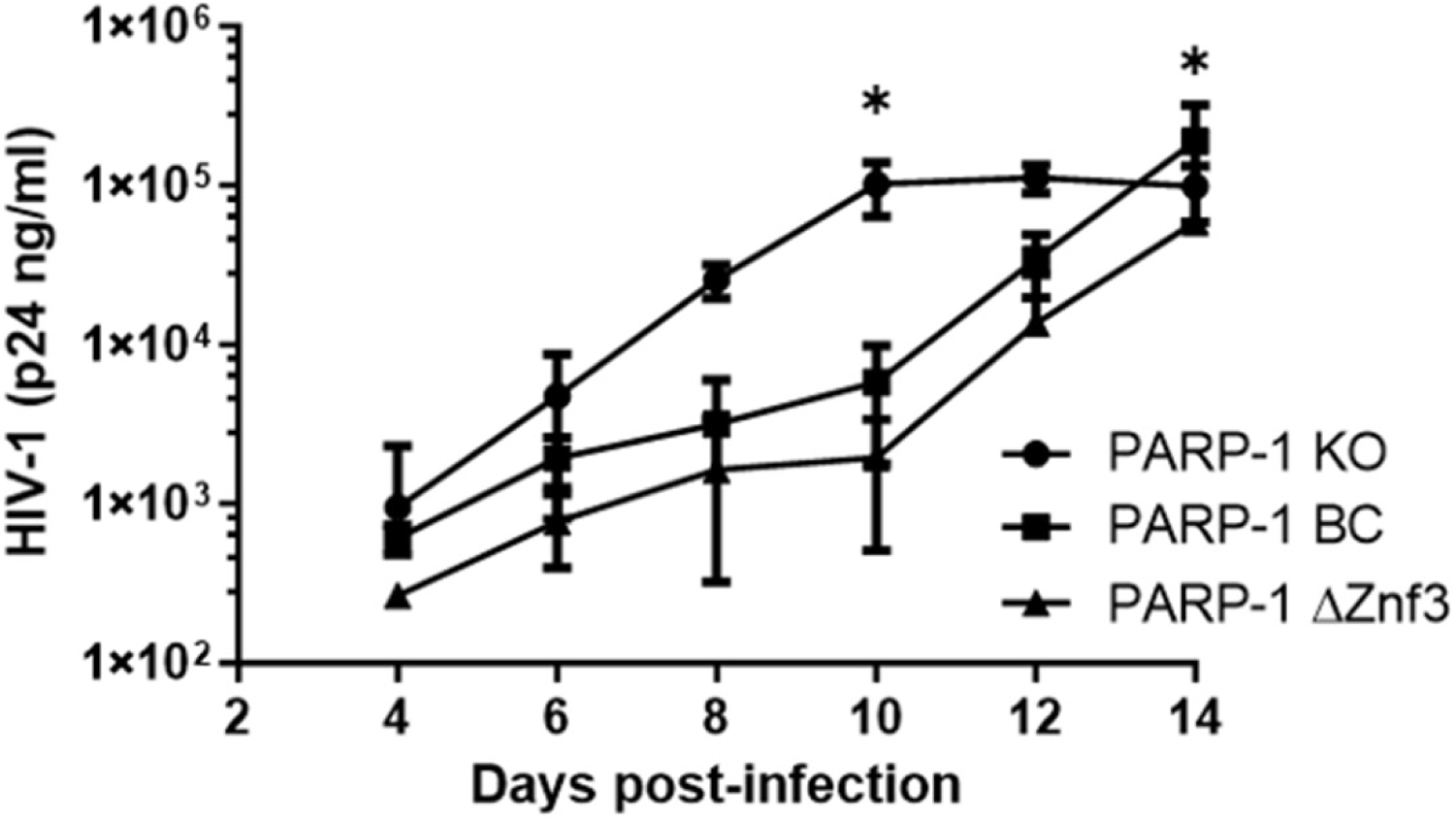
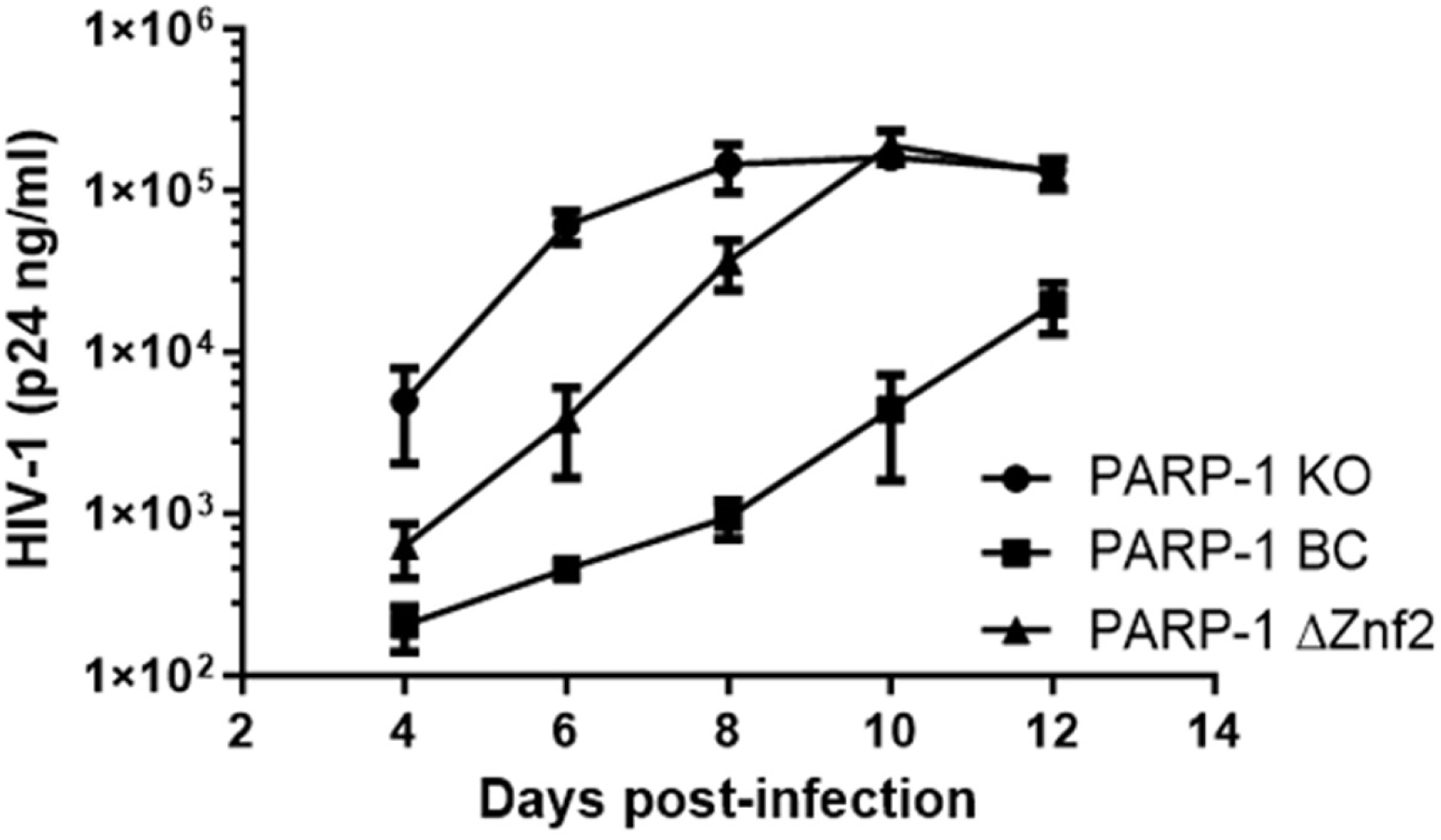
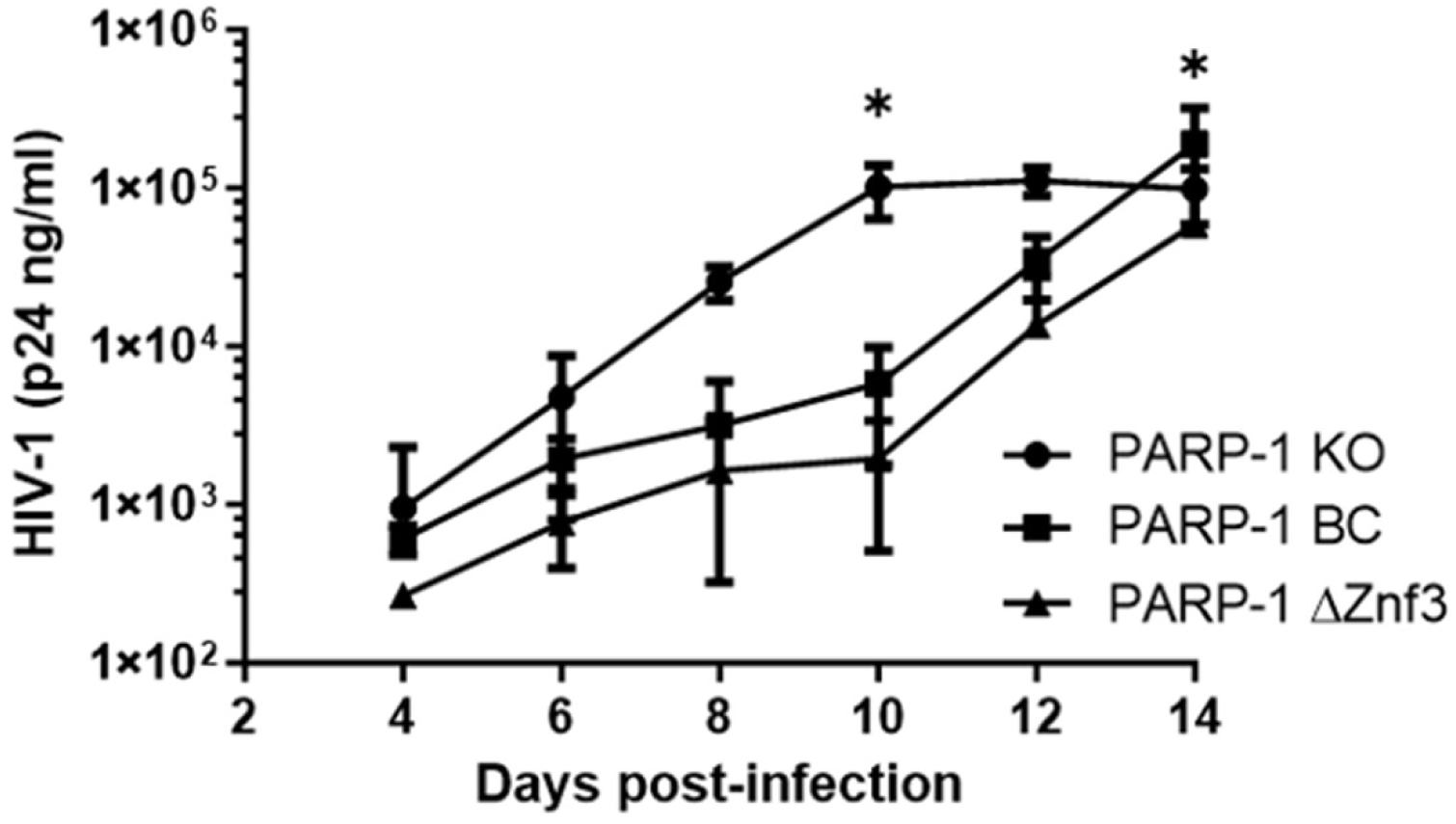
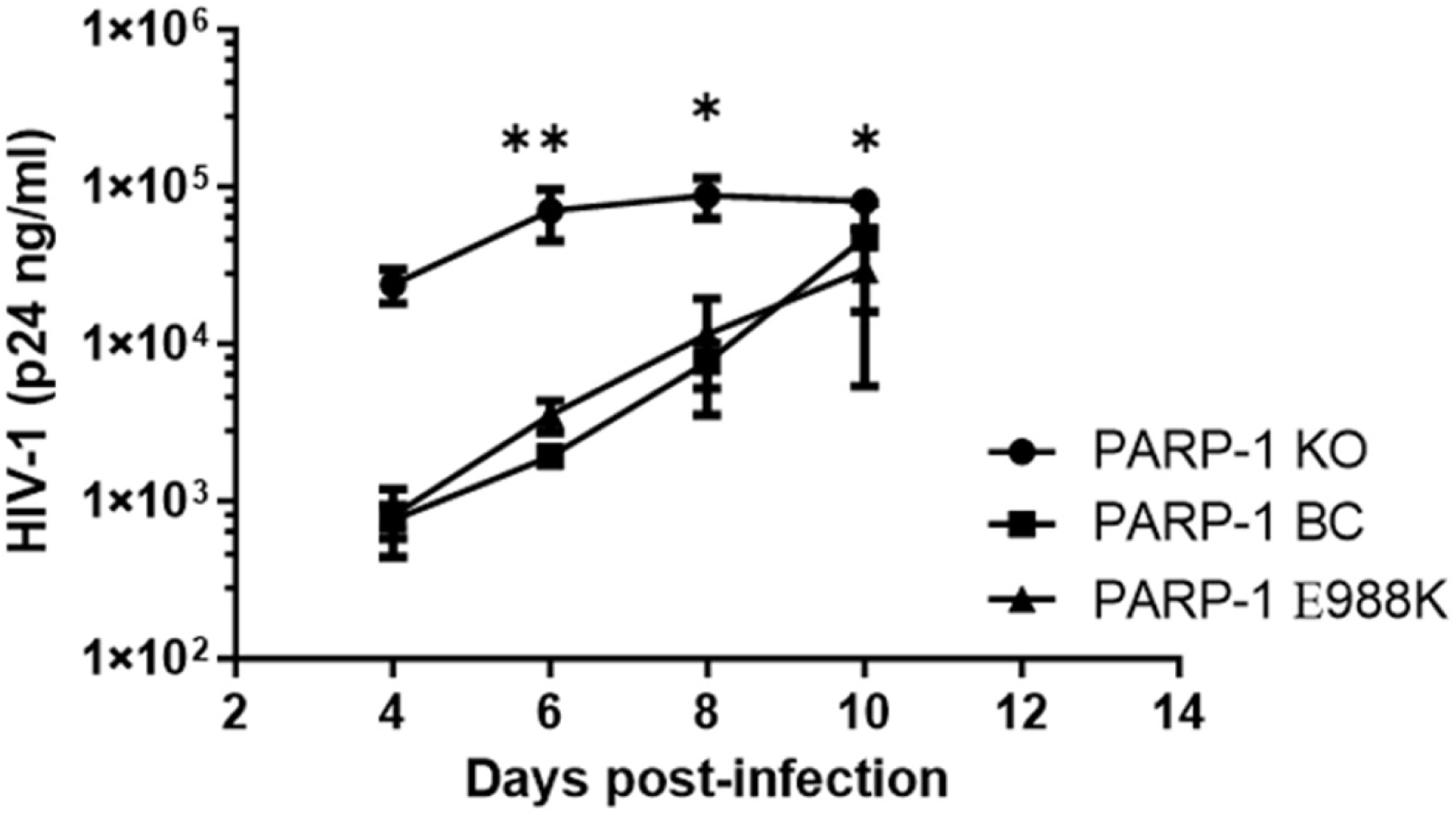
The DNA-binding domain of PARP-1 is required for the anti-HIV-1 function. (**a**) PARP-1 expression in PARP-1 KO SUP-T1 cells backcomplemented with wild-type or mutants PARP-1 lacking the zinc finger motifs 1, 2, and 3 (**a-I**) or carrying the point mutation E988K (**a-II**). (**b**) HIV-1_NL4-3_ replication in cells characterized in panel (**a**). Statistical analysis of the differences in HIV-1 replication in PARP-1 KO and cells backcomplemented with PARP-1 mutants was conducted in (**b**) by a 2-way ANOVA with Sidak’s post-hoc test. p > 0.05 (*) and > 0.01 (**). Non-significant differences (p </= 0.05) are not indicated.

DNA binding increases by 1000-fold the poly-ADP-ribosylation activity of PARP-1 (32). Therefore, to further evaluate the role of PARP-1 enzymatic activity in the anti-HIV-1 effect, we also replicated HIV-1_NL4-3_ in PARP-1-KO SUP-T1 cells backcomplemented with PARP-1 catalytic mutant E988K (**Fig. 5a-II**). This conserved residue is part of the catalytic triad, and this mutant cannot produce poly-ADP-ribosylation (33, 34). In agreement with the results obtained with PARP-1 inhibitors, PARP-1 E988K was fully active against HIV-1 (**Fig. 5b-IV**).

### PARP-1 deficiency increases Env levels in HIV-1 virions

Results described above suggest that PARP-1 antagonism enhances HIV-1 replication by increasing viral infectivity in an Env-dependent manner. Therefore, we evaluated the effect of PARP-1 on Env cellular production, processing, and incorporation into the virions. HIV-1_NL4-3_ was produced in HEK293T PARP-1-KO and -BC cells, concentrated by ultracentrifugation through a sucrose cushion, and the levels of gp160, gp120, and gp41 were determined by immunoblot in p24-normalized virions and in the producer cells. In both, virions and producer cells, the anti-gp41 antibody yielded strong signals and was able to detect gp160 (unprocessed Env) in addition to gp41 (processed Env). In contrast, the anti-gp120 antibody was poorly reactive, and although recognized gp120 in the producer cells, it failed to detect gp120 in the virions. This antibody also failed to recognize gp160 in the producer cells or the virions.

Using this approach, we found that PARP-1-KO virions harbored higher levels of gp41 and lower levels of gp160 than PARP-1-BC virions (**Fig. 6a-I**). However, no differences in the levels of gp41, gp120, gp160, or Gag were observed in HEK293T PARP1-KO or -BC producer cells (**Fig. 6a-II**). Densitometry analysis of multiple immunoblots, demonstrated that gp41 was 2 +/- 0.4 folds more abundant whereas gp160 was 3.3 +/- 0.5 folds less abundant in virions produced in PARP-1-KO cells than in PARP-1-BC cells (**Fig. 6b**). These findings suggest that PARP-1 deficiency enhances processed Env virion incorporation rather than Env production or processing in the producer cells, providing a potential explanation for the increased infectivity of virions produced in PARP-1-KO cells.

**Figure 6.**
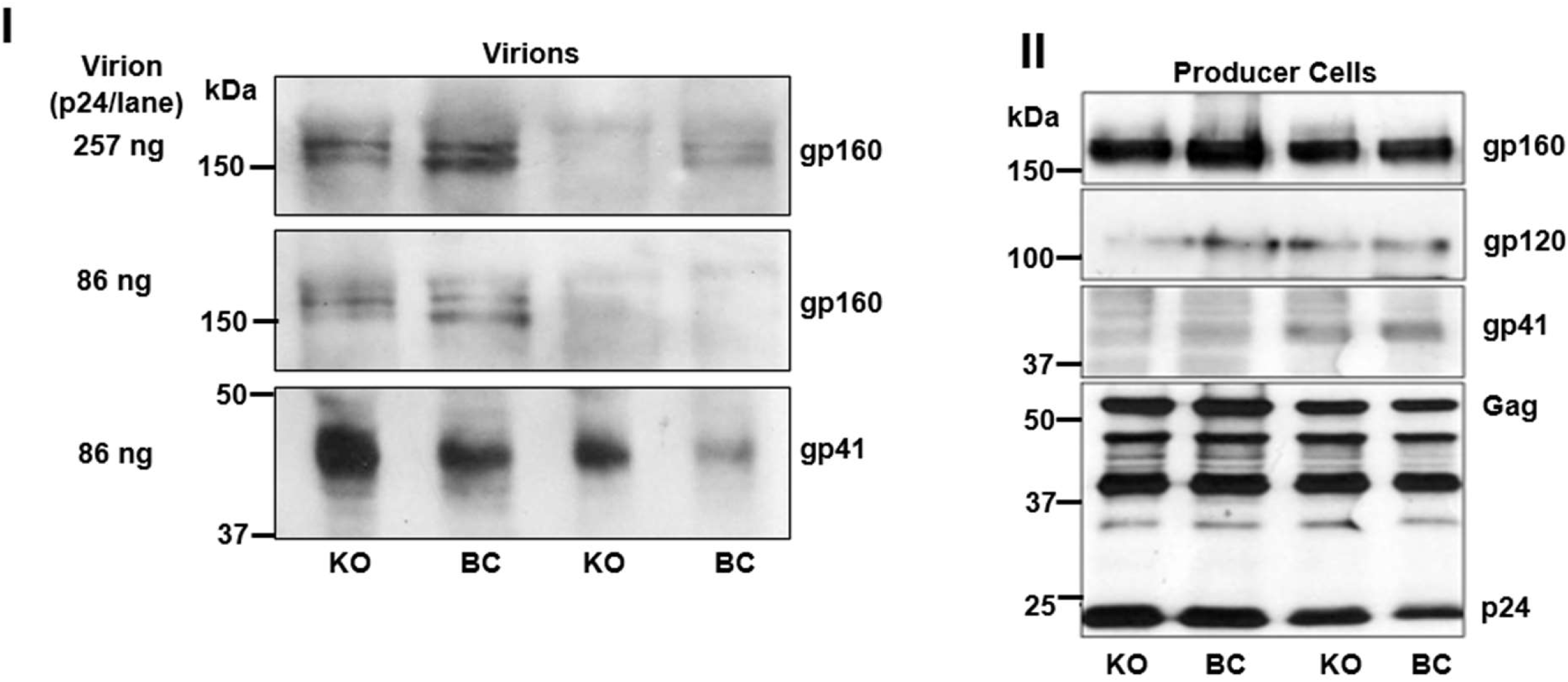
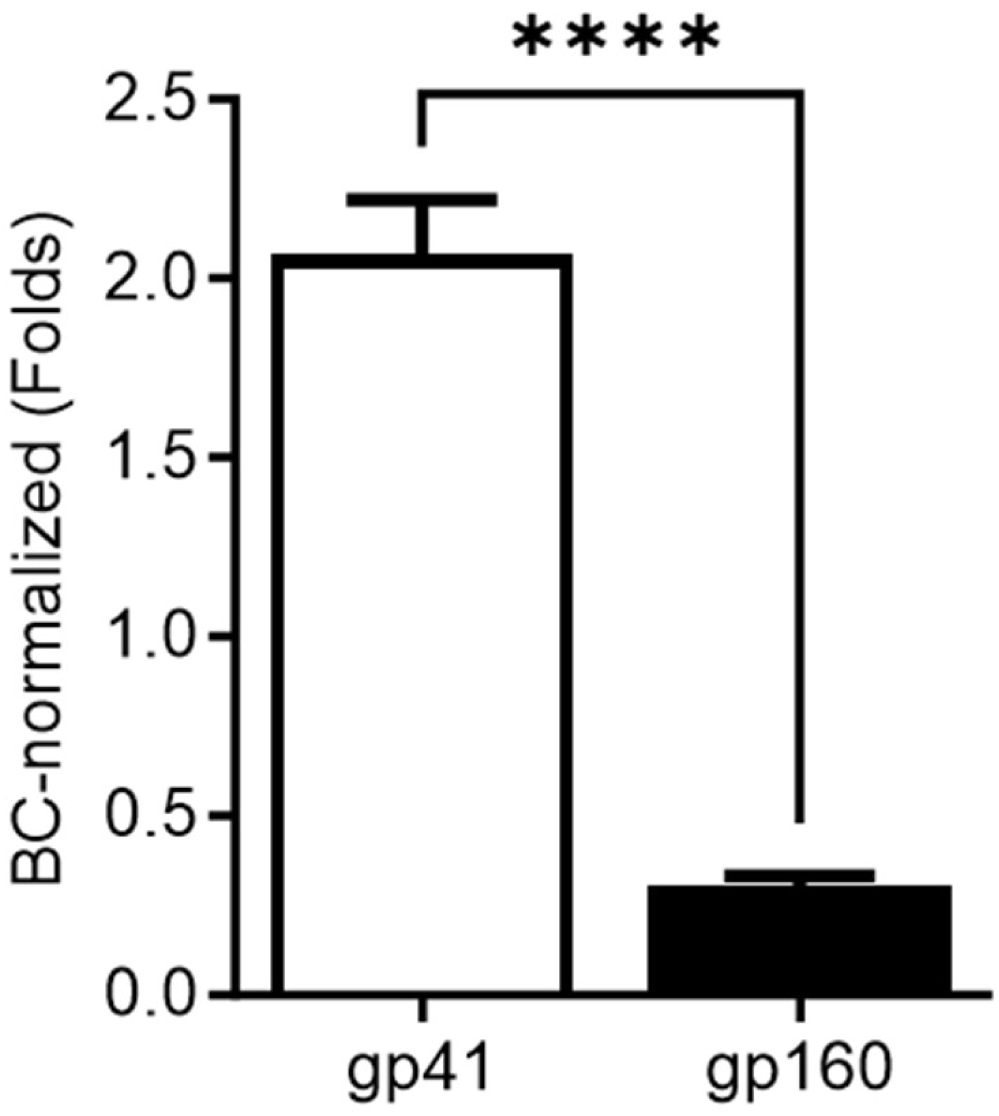
Effect of PARP-1 on Env expression and processing. (**a**) Levels of Env precursor (gp160) and processed forms (gp120 and gp41) in sucrose-cushion purified virions and their corresponding producer cells as detected by immunoblot with specific antibodies. HIV-1 Gag was also detected in the producer cells by immunoblot analysis. Results correspond to two independent experiments done in a single replicate. (**b**) Densitometry analysis of anti-gp41 immunoblots. Samples from the two viral preps of HIV-1_NL4-3_ KO and BC studied were evaluated in duplicate in three independent immunoblots using different amounts of HIV-1 p24 (257-13 ng per lane). In (**b**), statistical analysis was performed by a two-tailed t-test. p >0.0001 (****).

## DISCUSSION

Our data indicate that PARP-1 inhibition or deficiency (referred to as antagonism thereafter) enhances HIV-1 replication in human CD4+ T cells, although PARP-1 antagonism does not increase the susceptibility of the target cell to HIV-1 infection. This enhancing effect is Env-dependent, since only viruses entering through Env were affected. Indeed, infectivity of VSV-G-pseudotyped viruses is rather decreased than enhanced by PARP-1 antagonism. Wild-type HIV-1 produced in the absence of PARP-1 was more infectious in single-round infections in TZM-bl cells as well as in multiple-round infections in SUP-T1 cells, despite both cell lines express PARP-1. Therefore, PARP-1 antagonism enhances viral infectivity in the producer rather than in the target cell. In line with these observations, PARP-1 deficiency increased the levels of functional Env (Gp41) in the virions while decreased the amount of the inactive Env precursor protein, Gp160. The effect of PARP-1 on Env seems to be at the virion incorporation step because the amount or processing of Env in the producer cells were not affected by PARP-1 levels. Therefore, PARP-1 antagonism could enhance infectivity by this mechanism. Due to the large abundance of PARP-1 in the cell types relevant to HIV-1 infection *in vivo*, PARP-1 could contribute to the notoriously low abundance of Env in HIV-1 virions (35–37).

Pharmacologic inhibition of PARP-1 also enhanced HIV-1 infectivity and replication in an Env-dependent manner. However, this effect was only observed with the zinc ejector, INH2BP, that targets the DNA-binding domain of the enzyme, and no with inhibitors binding to the catalytic site. Phenocopying the effect of PARP-1 deficiency, HIV-1 produced in cells treated with INH2BP were more infectious in TZM-bl cells, but this enhancing effect was not observable with VSV-G-pseudotyped viruses. In addition, INH2BP did not increase HIV-1 provirus formation in SUP-T1 cells, suggesting an effect at a post-integration step of the viral life cycle. PARP-1 mutagenesis brought further support to the pharmacological studies indicating that the DNA-binding domain but not the poly-ADP-ribosylation activity of PARP-1 is required for the anti-HIV-1 effect.

HIV-1 infection activates caspase 3 and 7 via Env and Nef signaling in CD4 T cells (38–42). Activated caspase 3 and 7 cleave PARP-1, generating 89-kDa and 24-kDa fragments. The larger fragment lacks the DNA-binding domain and is released from the nucleus to the cytoplasm. In contrast, the 24-kDa fragment retains the DNA binding domain and persists irreversibly bound to cellular DNA, acting as a transdominant inhibitory mutant of PARP-1 functions (43–45). Therefore, our results predict that activation of these executioner caspases in response to infection will enhance Env virion incorporation through PARP-1 cleavage. This mechanism could explain, at least in part, why treatment of HIV-1 infected CD4 T cells with pan-caspase inhibitors decreased the production of infectious HIV-1 (39).

Furthermore, our findings indicate a differential role of PARP-1 in HIV-1 replication in the lymphocyte and myeloid compartments. While we found PARP-1 to impair HIV-1 replication in CD4 T cells, this enzyme was reported to promote HIV-1 replication in monocytes/macrophages (7, 10). This differential role of PARP-1 could be due to the differences in the virion Env incorporation mechanism in CD4 T cells and monocytes/macrophages. It is well stablished that there are cell type-specific differences in Env incorporation(46, 47). For example, it has been found that Vpr, which is dispensable for efficient replication of HIV-1 in CD4 T cells, promotes Env processing in monocytes/macrophages, increasing Env virion incorporation and infectivity (48, 49).

Our data also demonstrate that a proposed inhibitory role of PARP-1 in LTR/Tat-driven transcription (12, 28, 50, 51) is irrelevant in HIV-1-infected CD4 T cells. It was previously reported (12) that transfection of plasmids expressing PARP-1 and a LTR-driven reporter in non-infected HeLa cells engineered to express HIV-1 Tat, resulted in inhibition of the LTR/Tat-driven reporter expression. This inhibitory activity was proposed to be determined by the ability of PARP-1 to bind the HIV-1 trans-activation-responsive RNA and displace Tat·P-TEFb, that was observed in *in vitro* acellular studies (12, 28, 50, 51). However, we did not detect any increase in LTR/Tat-driven reporter expression in SUP-T1 cells lacking PARP-1 or treated with PARP-1 inhibitors following infection with VSV-G pseudotyped, single-round HIV-1. Furthermore, the high abundance of PARP-1 in HIV-1 susceptible cells (for example, figure 1a), indirectly support our view of a negligible inhibitory role of PARP-1 in LTR/Tat-driven transcription.

PARP-1 is a nuclear protein, and virion Env incorporation occurs at the plasma membrane; therefore, the effect of PARP-1 on virion Env incorporation is most likely indirect. PARP-1 regulates gene expression extensively (52–54), through catalytic-dependent and - independent mechanisms (55, 56). The later mechanism requires the binding of PARP-1 to the nucleosomal DNA at gene promoters and its interaction with other transcriptional regulators (2, 29, 57–59). Therefore, we propose that PARP-1 regulates the expression of host factors implicated in Env virion incorporation.

In conclusion, our data indicate that PARP-1 antagonism enhances HIV-1 infectivity, likely by increasing Env virion incorporation.

## MATERIALS AND METHODS

### Generation of PARP-1 deficient and control cell lines

SUP-T1 and HEK293T cells (kindly provided by Poeschla lab, Mayo Clinic) were used to derive PARP-1-knockout (KO) cells. These cell lines were generated by the transduction with lentiviruses expressing the 2 subunits of a PARP-1-specific zinc-finger nuclease (Sigma, *CKOZFN1116*). Transduced cells were subjected to single cell-cloning, and KO clones were selected and engineered to re-express wild-type PARP-1 [backcomplemented, (BC) cells] or PARP-1 mutants using the MLV-derived vector previously described (19).

### Generation of PARP-1 mutants

The Phusion Site-Directed Mutagenesis Kit (Thermo Fisher Scientific F541) was used. Zinc finger (ZnF)1 (aa 10-89) was deleted with the reverse primer (CH1) 5’-TCGATAGAGCTTATCCGAAGACTCC and forward primer (CH2) 5’-GAAGCTGGAGGAGTGACAGGCAAAGGCC, ZnF2 (aa 114-199) was removed with reverse primer (CH3) 5’-TGCAAAGTCACCCAGAGTCTTCTC and forward primer (CH4) 5’-CCAGGAGTCAAGAGTGAAGGAAAG, and ZnF3 (aa 234-373) was eliminated with reverse primer (CH5) 5-AAGCTTACTATCCTTGTCTTTTTC and forward primer (CH6) 5’-GCCTCGGCTCCTGCTGCTGTGAACTCC. The single point mutant E988K (<u>G</u>AG to <u>A</u>AG) was generated with reverse primer DG36 5-Tgttatatagtagagaggtgtc (mutagenic primer, upper-case mutated nucleotide) and the forward primer DG37 5’-agtacattgtctatgatattgc. The coding region of all mutants was verified by overlapping DNA sequencing.

### Generation of retroviruses

The HIV-derived viral vectors expressing the PARP-1 specific zinc-finger nucleases, the MLV-derived viral vectors expressing wild-type and mutants PARP-1, the single-round reporter virus (Hluc), and HIV-1_NL4-3_ were produced by calcium phosphate transfection of 3×10^6^ cells / T75 flask of HEK293T cells with the corresponding plasmids. For viral vectors expressing the zinc-finger nucleases, cells were transfected with pTRIP (60) expressing each of the PARP-1 zinc-finger nucleases (15 ug), the HIV-1 packaging plasmid pCMVΔR8.91 (15 ug), and the plasmid pMD.G (5 ug) encoding the Vesicular Stomatitis Virus glycoprotein G, as described in (61). pTRIP PARP-1 zinc-finger nucleases were generated by subcloning the PARP-1 zinc-finger nucleases obtained from pZFN1 and pZFN2 (Sigma, *CKOZFN1116*) in pTRIP at the NdeI and XhoI sites. This subcloning positioned the PARP-1 zinc-finger nucleases under the transcriptional control of the internal human cytomegalovirus (CMV) immediate-early promoter in pTRIP. The nucleases target the Zinc Finger 1 domain of PARP-1 at the target sequence CACTCCATCCGGCAC**cctgac**GTTGAGGTGGATGGG, where the cutting site is shown in bold lower-case letters. Plasmids pJZ308 (19) expressing PARP-1 (15 ug) and pMD.G (5 ug) were used to generate MLV-derived viruses. HIV-1_NL4-3_ was produced by transfection of a plasmid expressing the wild-type HIV-1 molecular clone NL4-3 that was obtained from the AIDS Research and Reference Reagent Program (catalog # 114). The replication-defective HIV-1 reporter virus (Hluc), which expresses LTR-driven luciferase from the NEF slot and contains a large deletion in ENV (19), was produced by transfection of pHluc (15 ug) and pMD.G (5 ug). HIV-1_NL4-3ΔEnv_ and HIV-1_NL4-3ΔEnv/VPR/NEF_ were produced by transfection of the corresponding expression plasmids (15 ug) and pMD.G (5 ug). HIV-1_NL4-3ΔEnv_ was derived by replacing the ENV gene in NL4-3 with the Env deleted mutant present in Hluc using NheI and SalI.

### Analysis of HIV-1 replication

SUP-T1 and SUP-T1-derived cells were plated at 0.25 x 10^6^ cells in 2 ml of culture medium in a T25 flask. HIV-1 was added to these cultures alone, in the case of experiments with PARP-1-deficient cells and their respective controls, or immediately after adding different treatments such as PARP inhibitors (3-Aminobenzamide, 3-ABA, 5 mM; NU 1025, 0.5 mM; and 5-Iodo-6-amino-1,2-benzopyrone, INH2BP, 50 uM), or DMSO. The flasks were kept in an upright position for 18 hrs and then the input virus and treatments were removed by extensive washing. Cells were spun down to remove the infectious medium and then the pellet was washed twice by centrifugation with 10 ml of culture medium each time. Then the cells were plated in a T25 flask in 4 ml of culture medium, and from day 4 post-infection were sampled every other day for HIV-1 p24 ELISA until peak viremia occurred, approximately 2 weeks after infection. In most of the experiments reported here, INH2BP 50 uM was added back to the cells after removing the input virus.

Viral replication was tracked by measuring HIV-1 p24 levels in the supernatant with an ELISA (ZeptoMetrix Corporation, catalog number 0801008) following manufacturer instructions.

Stocks of uninfected cells were kept for less than 3 weeks in culture, and compounds were kept at -20 °C. Doses of HIV-1 that produced approximately 0.1-1ug/ml of p24 by day 7-9 post-infection were used. These doses produced syncytia in SUP-T1 cells by day 7-9 post-infection. Importantly, we noticed that at higher viral doses, the effect of PARP-1 was less noticeable. Therefore, we used variable amounts of HIV-1 p24 in infection experiments to adjust for the differential infectivity of the multiple HIV-1 lots used.

### Single-round infection analysis

SUP-T1 parental and -derived cell lines and TZM-bl cells were plated at 0.1X10^6^ cells in 500 ul of culture medium per well in 24-well plates and infected with the non-replicating viruses. HIV-1 p24 doses similar to those used in HIV-1 replication experiments were used in these assays. In some experiments, cells were treated with INH2BP (50 uM) or DMSO right before infection, and input virus and compounds were removed 18 hrs later by extensive washing. Cells infected with viruses expressing luciferase were harvested 4 days after infection and analyzed for luciferase activity with a luminescence kit (Bright-Glo™ Luciferase Assay System, Promega, E2620) as described in (61).

### Quantification of HIV-1 provirus in infected cells

SUP-T1 cells were infected in the presence of INHB2 (50 uM) or DMSO, with doses of HIV-1 known to evidence the effect of PARP-1 in HIV-1 replication, and in the presence of the HIV protease inhibitor Indinavir (200 nM) to limit HIV-1 replication to a single cycle. Input virus was treated with turbo DNase before infection, as described in (19). Then, cells were washed 18 hrs after viral challenge, and cultured in the presence of Indinavir (200 nM). Analysis of the infected cells was done 2 weeks after infection to allow the non-integrated HIV-1 DNA to decay. Genomic DNA was purified from 10^6^ infected cells using the High pure PCR template preparation kit (Roche) and subjected to quantitative real-time PCR (qPCR). Extracted DNA (20 ng) was used for the detection of total HIV-1 cDNA (a segment of Gag), Alu-LTR junctions, and mitochondrial DNA by real-time PCR with primers and conditions previously described (19, 61). Total HIV-1 cDNA and Alu-LTR CT values were normalized to mitochondrial DNA levels, and fold changes were calculated using the ΔCt method. Additionally, genomic DNA extracted from SUP-T1 infected cells from five independent experiments were evaluated by Southern blot with a ^32^P-labeled probe that hybridizes within the LTR fragment obtained in the nested PRC step of the Alu-LTR junction PCR. For the southern blot, samples were normalized for their mitochondrial DNA content as determined by qPCR.

### Poly-ADP-ribose (PAR) ELISA and cell viability quantification

Protein-coupled or - uncoupled PAR was measured in cell lysates by ELISA (Catalog # 4520-096-K; Trevigen). SUP-T1 cells were treated for different amounts of time with DMSO or several PARP inhibitors and then washed in 1 ml of ice-cold 1X PBS by centrifugation at 1,000 x g for 6 min. The pellet was resuspended in 100 μl of cell lysis buffer supplemented with protease inhibitors and incubated on ice for 15 min with periodic vortexing. Then, SDS was added to the samples to achieve a final concentration of 1% and incubated at 100°C for 5 min. Once the cell extracts cooled to room temperature, 0.01 volume of Magnesium Cation (catalog # 4520-096-12; Trevigen) and 2 μl of DNAse I (2 Units/μl, catalog # 4520-096-07; Trevigen) were added, followed by incubation at 37°C for 90 min. To remove cellular debris, cell extracts were centrifuged 10,000 x g for 10 min at room temperature, and then, without disturbing the pellet, the supernatants (90 μl) were collected in a fresh tube for analysis. ELISA was performed following the manufacturer’s instructions. The viability of the cells treated with PARP-1 inhibitors was determined by measuring intracellular ATP. Cells were lysed in PBS-1% Triton, and ATP levels were measured with the CellTiter-Glo® Luminescent Cell Viability Assay (Promega, G7570) following the manufacturer’s instruction.

### Cell surface CD4 and CXCR4 analysis

PBS-washed cells (10^5^) were incubated on ice for 30 min in the dark in 50 ul of PBS containing 2 ul of APC-labeled mouse anti-human CD4 or anti-human CD184 (CXCR4) antibody (BD biosciences 561840 and 560936, respectively). Then, stained cells were analyzed by flow cytometry.

### Immunoblot analysis

PBS-washed SUP-T1 or HEK293T cells (3×10^6^) were lysed in 2X Laemmli sample buffer, and 100 ug of proteins per electrophoresis lane were analyzed. Sucrose cushion-purified virions were lysed in 6X Laemmli sample buffer and between 257 ng to 13.6 ng HIV-1 p24 were analyzed per lane. Proteins were resolved by SDS-PAGE and transferred overnight to PDVF membranes at 100 mA at 4°C. The membranes were blocked in TBS (25 mM Tris·HCl, 150 mM NaCl, pH 7.6) containing 10% milk for one hour and then incubated overnight at 4°C with anti-PARP-1 mouse monoclonal antibody C2-10 (Santa Cruz Biotechnologies sc-53643, 1/500) or a rabbit polyclonal IgG (Santa Cruz Biotechnologies sc-7150 1/1000). α-tubulin was detected as a loading control with the mouse monoclonal antibody B-5-1-2 (Sigma, T5168, 1/4000) for 2 hrs at 25°C. gp41 and gp160 were detected with a human antibody (1/500), Gp120 with goat polyclonal antibody (1/500), and Gag with anti-N-terminal p24 mouse monoclonal antibody (1/500). These anti-HIV-1 antibodies were incubated overnight at 4°C. The membranes were washed in TBS-0.1% Tween 20, and bound primary antibodies were detected with goat anti-mouse Ig-HRP (Sigma, 1/2000), goat anti-rabbit Ig-HRP (Sigma, 1/4000), goat-anti human Igs-HRP (1/2500), and mouse anti-goat Igs-HRP (1/5000). Binding of secondary antibodies was detected by chemiluminescence. All the antibodies were diluted in TBS-5% milk-0.05% Tween 20.

### Data analysis

Densitometry analysis of immunoblots was performed with ImageJ (https://ij.imjoy.io/). Statistical analysis was conducted with GraphPad Prism 5.01. Results were represented in the figures as follow: p > 0.05 (*), > 0.01 (**), >0.001 (***), >0.0001 (****). Non-significant differences (p </= 0.05) were not indicated in the figures.

## Acknowledgments

This work was supported by the National Institutes of Health grant numbers SC1GM115240 to ML from the National Institute of Allergy and Infectious Diseases and 5U54MD007592 from the National Institute on Minority Health and Health Disparities. We thank the Biomolecule Analysis, Genomic Analysis and the Cellular Characterization and Biorepository Core Facilities for technical help. These core facilities are supported by a Research Centers in Minority Institutions program grant 5G12MD007592 to the Border Biomedical Research Center in UTEP from the National Institute on Minority Health and Health Disparities, a component of NIH.

We thank Luis Valdes, Daniel Reyes, Alejandra Piña, and Christopher Farmer (UTEP) for technical help, and Armando Varela (UTEP) for critically reviewing this manuscript. We also thank the AIDS Research and Reference Reagent Program for providing plasmid expressing HIV-1 molecular clones NL4-3.

